# Epigenetic clock and methylation studies in vervet monkeys

**DOI:** 10.1101/2020.09.09.289801

**Authors:** Anna J. Jasinska, Amin Haghani, Joseph A. Zoller, Caesar Z. Li, Adriana Arneson, Jason Ernst, Kylie Kavanagh, Matthew J Jorgensen, Julie A. Mattison, Kevin Wojta, Oi-Wa Choi, Joseph DeYoung, Xinmin Li, Andrew W. Rao, Giovanni Coppola, Nelson B. Freimer, Roger P. Woods, Steve Horvath

## Abstract

DNA methylation-based biomarkers of aging have been developed for many mammals but not yet for the vervet monkey (*Chlorocebus sabaeus*), which is a valuable non-human primate model for biomedical studies. We generated novel DNA methylation data from vervet cerebral cortex, blood, and liver using highly conserved mammalian CpGs represented on a custom array (HorvathMammalMethylChip40). We present six DNA methylation-based estimators of age: vervet multi-tissue epigenetic clock and tissue-specific clocks for brain cortex, blood, and liver. In addition, two dual species clocks (human-vervet clocks) for measuring chronological age and relative age, respectively. Relative age was defined as ratio of chronological age to maximum lifespan to address the species differences in maximum lifespan. The high accuracy of the human-vervet clocks demonstrates that epigenetic aging processes are evolutionary conserved in primates. When applying these vervet clocks to tissue samples from another primate species, rhesus macaque, we observed high age correlations but strong offsets. We characterized CpGs that correlate significantly with age in the vervet. CpG probes hypermethylated with age across tissues were located near the targets of Polycomb proteins SUZ12 and EED, and genes possessing the trimethylated H3K27 mark in their promoters.

The epigenetic clocks are expected to be useful for age estimation of wild-born animals and anti-aging studies in vervets.

## Introduction

Non-human primates (NHPs) are regarded as critical animal models used in biomedical research (Jasinska et al. 2013; Vallender & Miller 2013; Meyer & Hamel 2014; Estes et al. 2018; Rogers 2018; Jasinska 2019) and key reference species used for constructing a comparative framework essential for evolutionary biology studies (Martin 2003; Chatterjee et al. 2009). NHPs, as compared with rodents, more closely resemble humans in terms of lifespan, life history strategies, cognitive processes, immunological behaviors (Bjornson-Hooper et al. 2019), inflammatory responses (Seok et al. 2013), and other health characteristics that are relevant to aging processes (Finch & Austad 2012). Therefore, NHPs are invaluable models for studying the pathomechanisms of age-related diseases, developing novel anti-aging treatments, and performing preclinical testing of such therapies before translation to human subjects (Colman 2018). Therefore, specialized tools are needed for the assessment of aging processes in NHP models in the context of the environmental and genetic factors regulating natural aging, the pathogenesis of age-related conditions, and the development and testing of anti-aging therapies.

On the molecular level, the process of aging is associated with epigenetic DNA modifications, such as DNA methylation (DNAm) of cytosine residues within CpG dinucleotides (5-methyl-cytosine) across the genome. DNA methylation levels have been used to develop multi-tissue estimators of chronological age and mortality risk (Horvath 2013; Chen et al. 2016; Horvath & Raj 2018; Levine et al. 2018; Lu et al. 2019; Bell et al. 2019). Whereas physiological conditions (e.g., BMI and menopause), pathologies (e.g., cancers and neurodegenerative diseases), and environmental factors (e.g., diet, exercise, and HIV infection) can affect the trajectory of DNAm age (Horvath & Levine 2015; Levine et al. 2016; Zheng et al. 2016; Quach et al. 2017; Horvath & Raj 2018; Kresovich et al. 2019), the pace of DNAm age is a heritable genetic trait linked to several genomic regions (Lu et al. 2016; Lu et al. 2018; Gibson et al. 2019).

The vervet monkey (genus *Chlorocebus*) is an Old World monkey frequently used as a model in biomedical research (Jasinska et al. 2013; Jasinska 2019) particularly for complex chronic diseases, many of which either are associated with aging or aggravate the process of aging. Vervets exhibit some aspects of human aging, including neurodegeneration (Postupna et al. 2017; Kalinin et al. 2013; Chen et al. 2018; Latimer et al. 2019), reproductive senescence and menopause (Atkins et al. 2014). Vervets have also been used in studies of reproductive physiology and obesity (Kuokkanen et al. 2016), the effects of genes and diet on growth and obesity (Schmitt et al. 2018), cardiometabolic health (Voruganti et al. 2013), physiological and behavioral stress responses (Fairbanks, Jorgensen, et al. 2011; Fairbanks, Bailey, et al. 2011; Jasinska, Pandrea, et al. 2020), and multi-tissue genetic regulation of gene expression, including that in tissues involved in stress responses (Jasinska et al. 2017). Because it is a natural host of simian immunodeficiency virus, which typically does not progress to immunodeficiency upon infection, the vervet is an established model for AIDS research (Pandrea et al. 2006; Chahroudi et al. 2012; Ma et al. 2013; Ma et al. 2014). Whereas HIV infection in humans is associated with age acceleration (Horvath, Stein, et al. 2018; Horvath & Levine 2015), the links between the benign course of simian immunodeficiency virus infection and aging in the vervet remain unknown.

Vervets from diverse African populations and the bottlenecked founder populations in the Caribbean have been phylogenetically characterized, and together with the genetically characterized extended pedigree Caribbean-origin vervets in the Vervet Research Colony (VRC) at Wake Forest School of Medicine, are used for genetic, gene-phenotype, developmental, and infectious disease studies (Jasinska et al. 2012; Warren et al. 2015; Huang et al. 2015; Svardal et al. 2017; Turner et al. 2018; Jasinska et al. 2017; Jasinska, Rostamian, et al. 2020; Ramensky et al. 2019; Schmitt et al. 2020). To advance use of the vervet as a model for developmental and aging studies, and facilitate DNAm-based assessments of the age effects of various environmental exposures, including preclinical testing of anti-aging therapies, here we created a multi-tissue epigenetic age estimator for the vervet, which is based on the blood, liver, and brain prefrontal cortex. Given that chronological ages are difficult to assess in free ranging monkeys, the epigenetic clock can also enable objective age assessment in wild vervet populations.

## Results

To identify age-related CpGs and develop a multi-tissue epigenetic age predictor for vervet monkeys, we leveraged developmental tissue resources from the VRC vervets comprising animals representing the entire vervet lifespan, from neonates to senile individuals, with known chronological ages accurate to 1 day as detailed in **Table 1**. We characterized DNAm in three tissues: the peripheral blood (N=240, from 1 day to 25 years of age), a classical immune tissue that is available through minimally invasive sampling and is routinely used for biomarker studies (Jasinska et al. 2009; Jasinska et al. 2012); the liver (N=48, from 0 day to 21 years of age), a key metabolic organ; and a region of the prefrontal cortex in the brain corresponding to the Brodmann area 10 (N=48, from 0 day to 22 years of age), a subregion implicated in personality expression and executive function. We generated high quality DNAm profiles from these samples using 36,727 CpGs located at highly conserved regions in the primates represented on the HorvathMammalMethylChip40.

**Table 1.** Description of the data by tissue type.

| **Tissue** | **N** | **No.**<br>**Female** | **Mean**<br>**Age** | **Min.**<br>**Age** | **Max.**<br>**Age** |
| --- | --- | --- | --- | --- | --- |
| Blood | 144 | 100 | 10.2 | 0.0027 | 25 |
| Cortex | 48 | 25 | 3.15 | 0 | 22.9 |
| Liver | 48 | 28 | 2.81 | 0 | 21.8 |

N=Total number of tissues. Number of females. Age: mean, minimum and maximum.

### Samples cluster by tissue type

Unsupervised hierarchical clustering of tissue samples on the basis of all tested CpG sites revealed three distinct clusters, one for each tissue type (**Supplementary Figure 1**). The clusters from the peripheral tissues, blood, and liver, grouped together, whereas the brain cortex cluster was more distant. Within the liver cluster, the samples from animals older than 8.7 years (N=7 individuals) formed a separate subcluster, thus suggesting marked differences in DNAm profiles between fully adult individuals versus immature individuals and young adults in this organ. The observations in the vervets supported previous results in humans showing that extensive tissue-specific remodeling of DNAm patterns occurs in the liver during aging (Bacalini et al. 2019).

### Epigenetic clocks

We used these high quality DNAm data to construct different epigenetic clocks for vervet only and for both human and vervet. For the construction of the dual human-vervet clock, we used the DNAm data previously generated with the HorvathMammalMethylChip40 in 852 human samples representing 16 tissues from individuals 0 to 93 years old (Morgello et al. 2001; Kabacik et al. 2018; Horvath, Stein, et al. 2018). Our clocks for vervet monkeys can be distinguished along three dimensions (tissue type, species, and measure of age). We used a combined set of all samples to train a multi-tissue clock (pan-clock) suited for age predictions across different tissue types included in the clock construction. We also created clocks tailor-made for specific tissues/organs, which were trained on the basis of the samples from individual tissue types: the blood-clock, the liver-clock, and the brain cortex-clock. We anticipate that pan-clock may provide a proxy for tissues for which tissue-specific clocks are not available.

While the multi-tissue vervet clock applies only to vervets, we also created dual species clocks, referred to as human-vervet clocks, for estimates of chronological age and relative age. Relative age is the ratio of chronological age to maximum lifespan (i.e., the maximum age of death observed in the species). Thus, relative age takes on values between 0 and 1. The maximum lifespan observed for humans and vervets was 122.5 and 30.8 years, respectively. Relative age allows alignment and biologically meaningful comparison between species with different lifespan (vervet and human), which is not afforded by mere measurement of chronological age.

To arrive at unbiased estimates of the epigenetic clocks, we used leave-one-out (LOO) cross-validation of the training data. The cross-validation study reports unbiased estimates of the age correlation R (defined as Pearson correlation between the age estimate (DNAm age) and chronological age) as well as the median absolute error (mae) measuring the deviation between the predicted and observed age (for chronological age in years). As indicated by its name, the vervet multi-tissue clock is highly accurate in age estimation of the different tissue samples (R=0.98 and median error 0.89 years, **Figure 1A**). The multi-tissue clock also performs well when restricting the analysis to samples from a given tissue type: R=0.98 in the blood, R=0.99 in the liver and R= 0.91 in the brain cortex (**Figure 2**). We also developed highly accurate vervet clocks for single tissues: blood (R=0.98, **Figure 1B**), cerebral cortex (R=0.95, **Figure 1C**), and liver (R=0.99, **Figure 1D**). The accuracy of the multi-tissue clock for the vervet (r=0.98) exceeded the accuracy of the human pan-clock (r=0.96) (Horvath 2013) the accuracy of mouse multi-tissue clocks, which have been reported to range from r=0.79 to r=0.89 (Thompson et al. 2018).

**Figure 1:**
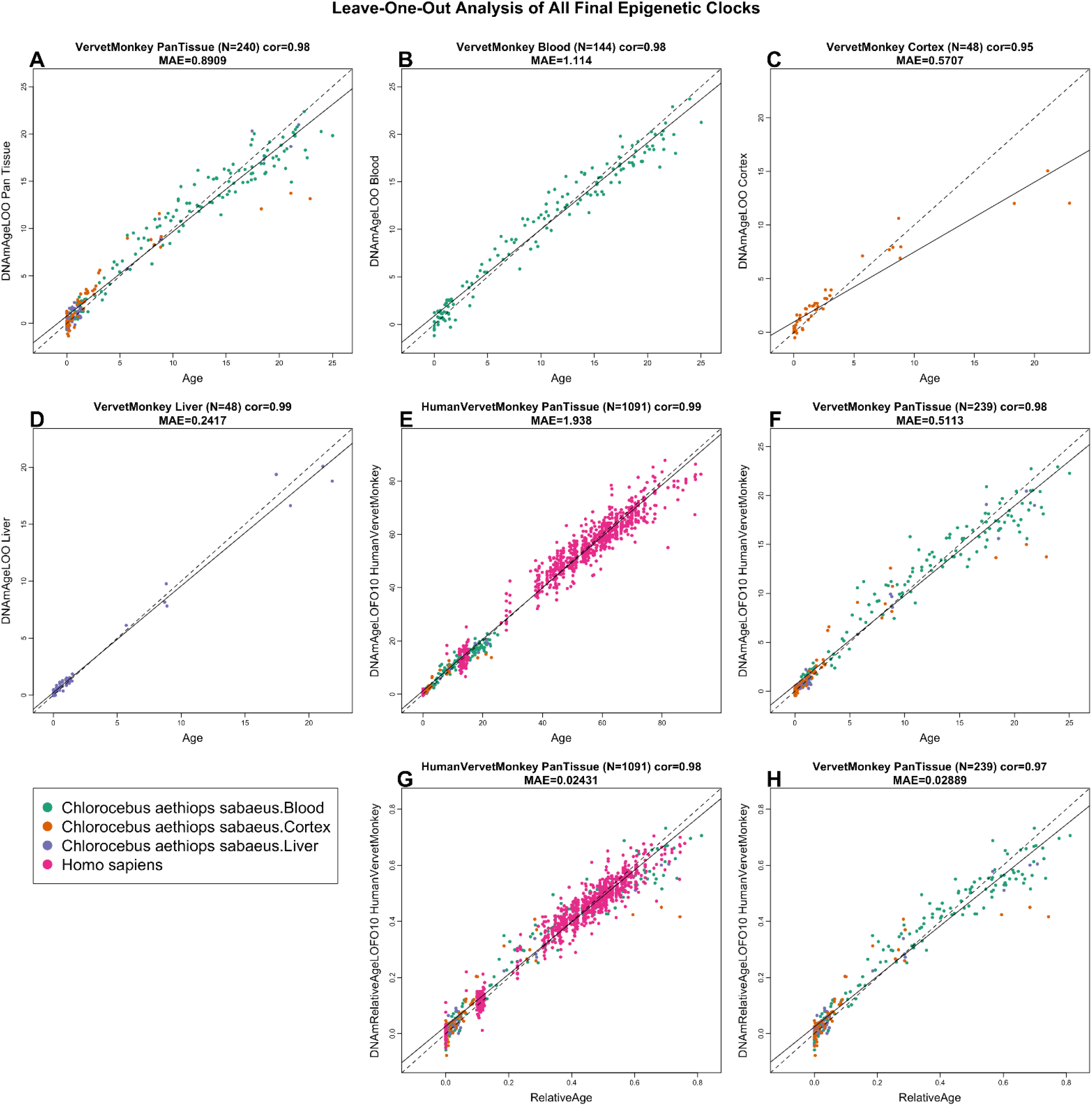
Cross-validation study of epigenetic clocks for vervet monkeys and humans. A-D) Four epigenetic clocks that only apply to vervet. Leave-one-sample-out estimate of DNA methylation age (y-axis, in units of years) versus chronological age for A) all available vervet tissues, B) vervet blood, C) vervet cerebral cortex, D) vervet liver. Ten fold cross validation analysis of the human-vervet monkey clocks for E,F) chronological age and G,H) relative age, respectively. E,G) Human samples are colored in magenta and vervet samples are colored by vervet tissue type, and analogous in F,H) but restricted to vervet samples (colored by vervet tissue type). Each panel reports the sample size (in parenthesis), correlation coefficient, median absolute error (MAE).

**Figure 2.**
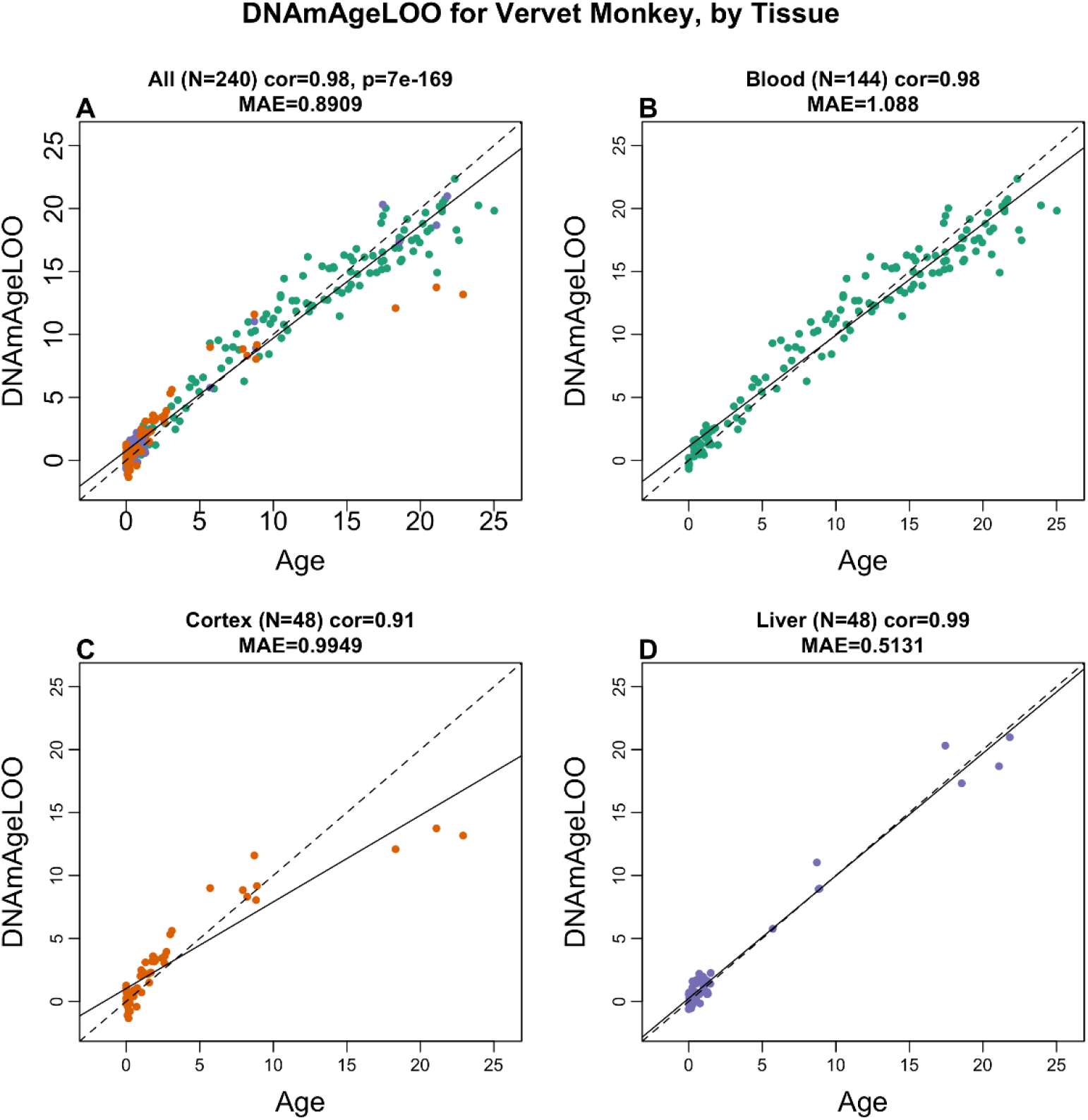
The multi-tissue epigenetic clock for vervets applied to individual tissues. Leave-one-sample-out estimate of age based on DNA methylation data (x-axis) versus chronological age (in units of years) for A) all tissues, B) blood, C) cerebral cortex, D) liver. Each panel reports the sample size, Pearson correlation coefficient and median absolute deviation (median error).

We developed two dual species clocks based on our vervet samples and previously characterized human tissues (Morgello et al. 2001; Horvath, Stein, et al. 2018; Kabacik et al. 2018). The human-vervet clock for chronological age (R=0.99 for the human and vervet samples and R=0.98 for the vervet samples, **Figure 1E,F**) and relative age (R=0.98 for the human and vervet samples and R=0.97 for the vervet samples, **Figure 1G,H**).

We tested the performance of the vervet blood clock in longitudinal blood samples from 14 individuals collected at two different time points. In these samples, we correlated the changes in DNAm age predicted on the basis of the vervet blood clock with the changes in the actual chronological age (**Supplementary Figure 2**). In all pairs of samples from the same animal, the samples collected later were correctly predicted to be from an older animal.

### Vervet clock applied to other primates

To determine the cross-tissue performance and the cross-species conservation of the vervet multi-tissue clock, we applied the vervet pan-clock to an array of tissues from key organs from two primate species: macaque (N=283 samples from eight tissues) and humans (N=852 from 16 tissues). The data from rhesus macaque are described in a companion paper (Horvath, Zoller, et al. 2020).

We observed an overall moderate to high correlation between the chronological age and predicted age based on the vervet multi-tissue clock: R=0.77 for macaque (**Supplementary Figure 3**) and R=0.62 for human (**Supplementary Figure 4**). The highest correlation in individual tissues was observed in blood (R=0.83 in macaque and R=0.81 in humans) and in skin (R=0.79 in macaque and R=0.9 in humans) (**Figure 3, Figure 4**). Strictly speaking, it is not possible to compare the correlation coefficients across the different tissues since it greatly depends on the underlying age distribution (e.g. minimum and maximum age) and to a lesser extent on the sample size. The vervet clock is poorly calibrated in other primates such as rhesus macaque and human as reflected by an “offset” that leads to a high median error in many tissues (**Figure 3, Figure 4**). However, the vervet clock leads to moderately high correlation coefficients in rhesus adipose (R=0.79), blood (R=0.83), brain cortex (R=0.75), liver (R=0.89), muscle (R=0.78) and skin (R=0.79) (**Figure 3**).

**Figure 3.**
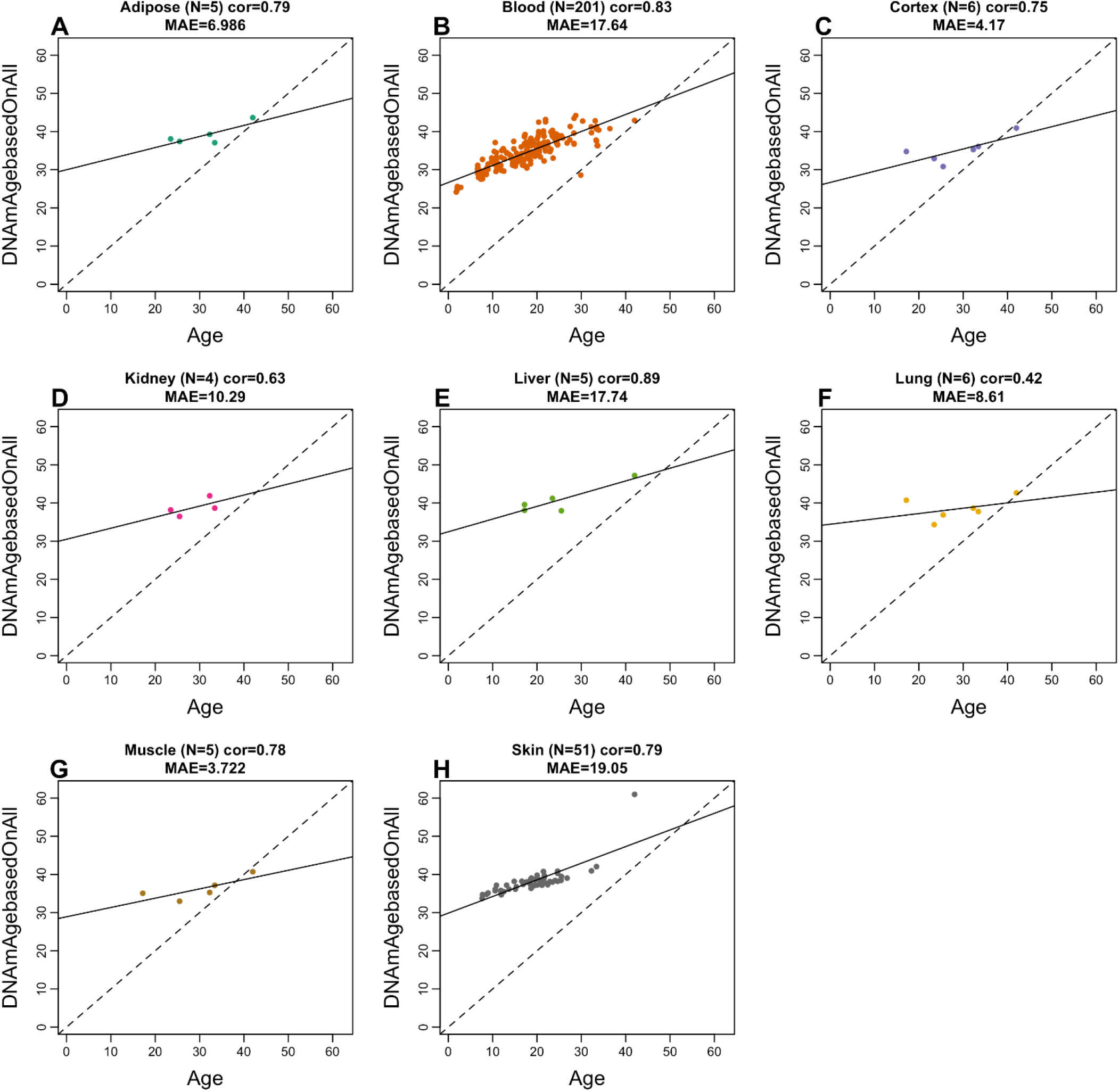
Multi-tissue vervet monkey clock applied to tissues from rhesus macaques. Each dot corresponds to a tissue sample from rhesus macaques: A) adipose, B) blood, C) brain cortex, D) kidney, E) liver, F) lung, G) muscle, H) skin. The y-axis reports the age estimate according to the multi-tissue vervet clocks. The predicted DNAm age in macaque tissues according to the vervet pan-clock (y-axis) and chronological age of the rhesus specimens (x-axis). The number of samples is shown in parentheses; cor – Pearson’s correlation, MAE – median absolute error.

**Figure 4.**
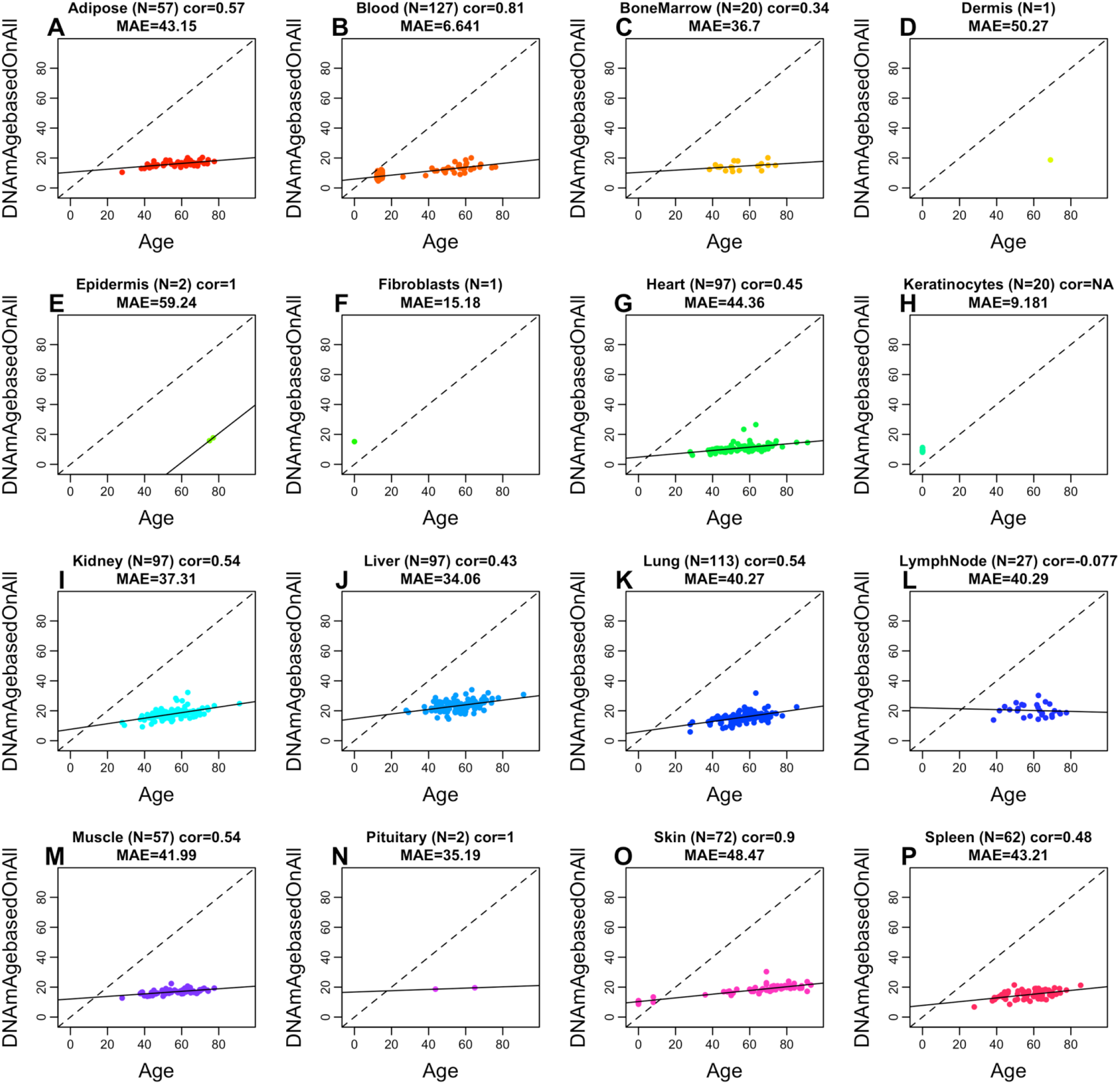
Multi-tissue vervet clock applied to 16 tissue types from humans. Each dot corresponds to a human tissue samples. The predicted DNAm age in human tissues according to the vervet multi-tissue clock (y-axis) and chronological age of the human specimens (x-axis). The number of samples is shown in parentheses; cor – Pearson’s correlation, MAE – median absolute error.

### Age related CpGs in vervets

In total, 36,727 probes from HorvathMammalMethylChip40 were aligned to specific loci proximate to 6,110 genes in the vervet monkey (*Chlorocebus_sabaeus.ChlSab1.1.100*) genome. The probes in this array were selected based on conservation in mammalian genomes, thus, our findings have high translatability into humans and other mammals. Epigenome-wide association studies (EWAS) of chronological age revealed that the age-related changes in DNAm are to a marked extent tissue-specific in the vervet monkey (**Figure 5A**).

**Figure 5.**
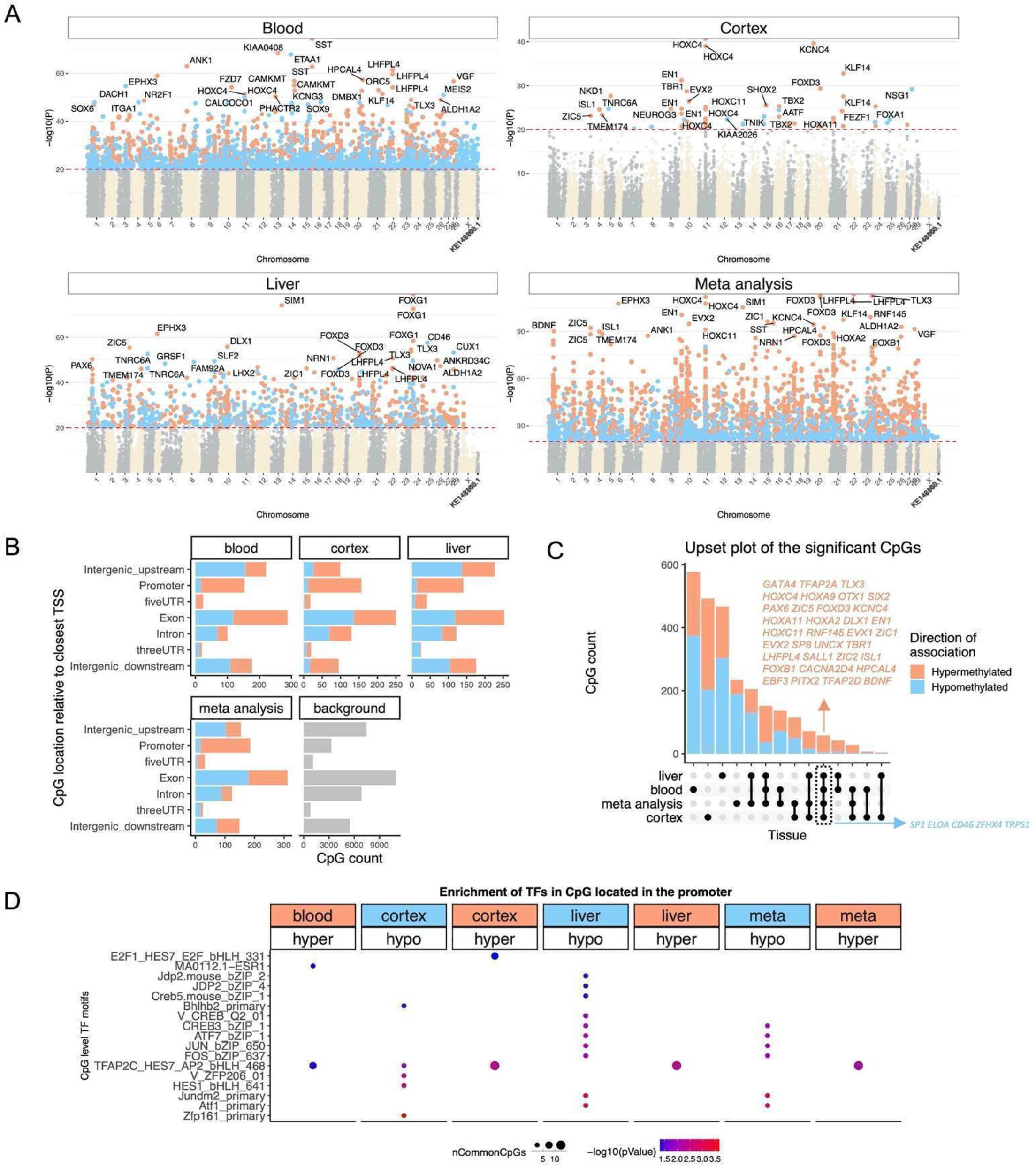
Epigenome-wide association study of age in tissues from Chlorocebus sabaeus. A) Manhattan plots of the EWAS results in different tissues. Stouffer meta analysis was used to combine the results across different tissues. The coordinates are estimated based on the alignment of Mammalian array probes to ChlSab1.1.100 genome assembly from ENSEMBL. The direction of associations with p < 1e-20 (red dotted line) is highlighted by red (hypermethylated) and blue (hypomethylated) colors. Top 30 CpGs was labeled by the neighboring genes. B) Location of top CpGs in each tissue relative to the closest transcriptional start site. Top CpGs were selected at p < 10^−10^ and further filtering based on z score of association with chronological age for up to 500 in a positive or negative direction. The number of selected CpGs: blood, 1000; cortex, 777; liver, 1000; meta-analysis, 1,000. The grey color in the last panel represents the location of 35,898 mammalian BeadChip array probes mapped to ChlSab1.1.100 genome. C) Upset plot representing the overlap of aging-associated CpGs based on meta-analysis or individual tissues. Neighboring genes of the overlapping CpGs were labeled in the figure. D) Transcriptional motif enrichment for the top CpGs in the promoter and 5’UTR of the neighboring genes. The motifs were predicted using the MEME motif discovery algorithm, and the enrichment was tested using a hypergeometric test (Bailey et al. 2009). In total, 19,087 CpGs were predicted to be located on the motifs and were used as the background. nCommonCpGs indicates the number of target CpGs that overlapped with the background CpGs on the analyzed motif.

Pairwise scatter plots of the age EWAS signals (**Supplementary Figure 5**) revealed a moderate positive correlation among the peripheral tissues, blood and liver (r=0.63), whereas the correlations among the peripheral tissues and the brain cortex were markedly lower, i.e., liver and cortex (r=0.219), and blood and cortex (r=0.14). However, the moderate to low conservation and differences in p-value ranges in our analyzed tissue types may reflect a moderate sample size in non-blood tissues.

The age EWAS results suggest that the cerebral cortex has the lowest number of DNAm changes associated with age (N= 916) compared to blood (N= 12,334) and liver (N= 4,454) at a nominal p-value < 10^−10^. This lower number of the age-associated DNAm alterations is probably not due to statistical differences because both brain and liver had the same number of samples (N=48) and with similar age distribution (**Table 1**). Rather, we hypothesize that it can be attributed to the post-mitotic state of neurons in the brain compared to blood and liver cells. However, the lower number of age related CpGs in the brain could reflect heterogeneity of cell types in the bulk cortical brain tissues or technical issues (Zeisel et al. 2015; Tasic et al. 2016) that are difficult to dissect with absolute consistency.

The genomic localization of top age-related DNAm changes and the proximate genes in each tissue are as follows (**Figure 5A**). The most significant EWAS signals in the blood are in a *KIAA0408* exon (z = 19.1) and in a promoter of the *SST* gene (z = 18.3), which is encoding somatostatin acting as a negative regulator of growth hormone slowing aging in humans (Bartke 2019). In the cerebral cortex, the strongest EWAS signals are in an exon of the *KCNC4* gene (z = 13.3) and in a promoter of the *HOXC4* gene (z =13.5), which is a homeobox gene crucial during erythroid lineage differentiation (Bhatlekar et al. 2018), and which expression is associated with age in bone marrow stromal cells (Pasumarthy et al. 2017) and skin cell differentiation and tumors (Rieger et al. 1994). Given the identification of a gene involved in erythroid lineage among the brain clock sites, it is pertinent to note that the brains were perfused to remove blood prior to dissection. The top EWAS signals in the liver are in a promoter of the *FOXG1* gene (z = 18.9), which mutations cause a severe neurodevelopmental disorder, the Rett syndrome (Ariani et al. 2008), and in an intron of the *SIM1* gene (z = 18.3).

In the meta-analysis across these three tissue types, the top age-related DNAm changes included hypermethylation in *LHFPL4* exon (z = 22.7), an exon of the *FOXD3 gene* (z = 22.6), which is a transcription repressor essential in embryogenesis controlling the multipotent mammalian neural crest, neuronal differentiation and fate (Teng et al. 2008; Mundell & Labosky 2011; Respuela et al. 2016), and a promoter of the *TLX3* gene (z = 22.6), which is a transcription factor acting as a master regulator of neuronal differentiation in embryonic development and in embryonic stem cells (Kondo et al. 2008; Xu et al. 2008).

The most significant enrichment of EWAS signals for biological terms according to the GREAT analysis was observed for the genes associated with the CpG probes hypermethylated with age in meta analysis (**Supplementary Figure 6**). The top enrichment was observed for the targets of the Polycomb proteins SUZ12 (BENPORATH_SUZ12_TARGETS) and EED (BENPORATH_EED_TARGETS), and genes possessing the trimethylated H3K27 mark in their promoters (BENPORATH_ES_WITH_H3K27ME3). Given that the polycomb complexes are involved in the chromatin remodeling resulting in epigenetic silencing of genes, such as homeobox genes, and that they were implicated in the modulation of brain aging (Kennerdell et al. 2018), the enrichment for the polycomb targets and genes with H3K27ME3 in their promoters among the hypermethylated CpGs with age is consistent with age-related gene repression. A highly significant enrichment was also observed for genes involved in DNA binding (sequence-specific DNA binding, DNA binding, sequence specific DNA binding transcription factor activity) suggesting the role of transcriptomic regulation in aging, and nervous system (nervous system phenotype, abnormal nervous system morphology) and development (lethality during fetal growth through weaning).

We analyzed the distribution of age-associated CpGs in different tissues across different genomic regions, including promoters, UTRs, exons, introns, and intergenic sequences (**Figure 5B**). The percentage of age-associated CpGs in the promoters was higher in all models (EWAS in individual tissues and in meta-analysis) compared to the background (i.e., all CpGs on the chip). Moreover, the most significant age-related DNAm changes were hypermethylation events in the promoters and 5’UTRs. These findings corroborate the role of CpGs in the promoters and the gain of methylation with age observed in other species (Johnson et al. 2012).

Using the upset plot analysis, we identified the CpGs that showed consistent age associated DNAm changes in multiple tissues (**Figure 5C**). We observed 58 of such shared CpGs undergoing age-related DNAm alterations in blood, cerebral cortex, liver, and meta-analysis. They were located proximate to 39 genes and included 34 CpGs hypermethylated and 5 CpGs hypomethylated with age. The top affected CpGs included hypermethylation in *LHPL4* exon, *FOXD3* exon, *TLX3* promoter, and hypomethylation in *SP1* exon, a CpG downstream of *CD46*, and *TRPS1* intron (**Figure 5C**). Some of these genes are implicated in aging phenotypes. For example, *SP1* is a key regulator of mTORC1/P70S6K/S6 signaling pathway (Astrinidis et al. 2010; Finotti et al. 2015) and is involved in several aging-associated diseases including cancer (Zhang et al. 2014), hypertension (Yang & Kaye 2009), atherosclerosis (Dunzendorfer et al. 2004), Alzheimer’s (Santpere et al. 2006), and Huntington diseases (Chen-Plotkin et al. 2006).

We examined the transcriptional factor motifs enriched for the top CpGs located in promoters or 5’UTRs with DNAm changes in either direction of each tissue (**Figure 5D**). The top TF motif most significantly enriched for the top EWAS CpGs was Zfp161 (ZBTB14 in human) motif hypomethylated with age in the cerebral cortex. At the next levels, hypomethylated CpGs showed a strong enrichment for the Atf1 and several immune-related TF motifs such as Jundm2, FOS, JUN, and CREB in the liver. TFAP2C was a TF motif hypermethylation in all tissues. This motif is involved in cell-cycle arrest, germ cell development, and it is implicated in several types of cancer (Bryant et al. 2012; Penna et al. 2013).

## Discussion

Based on the ~37 thousand CpG probes on the custom Mammalian methylation array (HorvathMammalMethylChip40), we generated DNA methylation data from three tissue types (brain, blood, liver) in the vervet monkey. These samples represent the most comprehensive dataset thus far of methylomes in vervets across multiple tissues and ages. We obtained high quality DNAm data, as reflected in the perfect clustering pattern of the samples by tissue type without any intermixture between different tissue samples within the clusters. Using these DNAm data, we trained and validated highly accurate age estimators (epigenetic clocks) that apply to the entire life course (from birth to old age), and identified genes associated with the aging process in the vervet.

These data allowed us to construct a highly accurate multi-tissue age estimator (pan-clock) based on three vervet tissue types (brain, blood, liver), and clocks developed based on individual vervet tissues. This gives us confidence that these vervet clocks will work on new vervet samples from other tissue types as well. However, we cannot rule out that these clocks could fail in some highly specialized cell types. Epigenetic age estimators that focus on specific tissues or cell types can have greater accuracy than multi-tissue age estimators (Horvath, Oshima, et al. 2018).

The vervet pan-clock showed an overall high to moderate accuracy of age estimates in a wide range of tissues in macaque and humans (except the lymph nodes in humans), despite the phylogenetic distance of ~12 Mln between the vervet and macaque, and ~29 Mln years between the vervet and human lineages (Kumar et al. 2017). The preservation of the predictive effect of the clock CpGs suggests a marked conservation of the aging mechanisms in the Catarrhini parvorder. Given that the vervet pan-clock can effectively predict age in numerous tissues, even in a different primate species, we anticipate that the pan-clock may also serve as an effective age estimator for various tissues beyond these included in the pan-clock construction.

Marked conservation of the clock constructed based on the Caribbean-origin vervets across different primate species suggest that this clock could provide accurate age estimates not only for different species but also for vervets from African populations, although we did not test it in these populations. The genetic architecture of VRC vervets is simplified compared to that of wild African vervets due to genetic bottlenecks (Warren et al. 2015). While the methods used to create the clock presumably favors methylation sites for which genetic variance is low, one concern is that the bottlenecks may have driven to fixation in the VRC certain genetic variants that might not be fixed in other populations and that might confound age estimates in wild Caribbean vervets and especially wild African vervets. Therefore, the vervet clock results should be interpreted with caution and would benefit from a validation in African vervet samples.

Epigenetic clocks for humans have found many biomedical applications including the measure of age in human clinical trials (Horvath & Raj 2018; Fahy et al. 2019). This instigated development of similar clocks for mammals such as mice (Petkovich et al. 2017; Cole et al. 2017; Wang et al. 2017; Stubbs et al. 2017; Thompson et al. 2018; Meer et al. 2018). While rodent models have obvious advantages, it can be challenging to translate findings from rodents to primates (Perrin 2014; Hatzipetros et al. 2014). NHPs play an indispensable role in aging studies and preclinical work of anti-aging treatments (Lankau et al. 2014; Mattison & Vaughan 2017). Lifespan and healthspan studies, as well as assessments of anti-aging interventions in primates remain costly and time consuming. The development of suitable biomarkers promises to greatly reduce the costs and time needed for carrying out studies in these primates. To increase the chance that findings in vervets translate to humans, we created dual species clocks, human-vervet clocks, for absolute and relative age. The bias due to differences in maximum lifespan is mitigated by the generation of the human-vervet clocks for *relative* age clock, which embeds the estimated age in context of the maximal lifespan recorded for the relevant species. The mathematical operation of generating a ratio also generates a more biologically meaningful value, because it indicates the relative biological age and possibly fitness of the organism in relation to its own species. The high accuracy of these clocks demonstrates that one can build epigenetic clocks for two species based on a single mathematical formula.

A critical step toward crossing the species barrier was the use of a mammalian DNA methylation array that profiled 37 thousand CpG probes that were highly conserved across numerous mammalian species. We expect that the availability of these clocks will provide a significant boost to the attractiveness of the vervet as a translational model for health, developmental, and aging research. The vervet pan-clock and tissue-specific clocks are biomarkers that can facilitate studies of the course of biological aging in the context of various genetic factors and environmental exposures (for example, preclinical testing of rejuvenating therapies).

Beyond their utility, these epigenetic clocks reveal several salient features with regards to the biology of aging. First, the vervet multi-tissue clock re-affirms the implication of the human multi-tissue clock, which is that aging might be a coordinated biological process that is harmonized throughout the body. Second, the ability to combine these two multi-tissue clocks into a single human-vervet multi-tissue clock attests to the high conservation of the aging process across two evolutionary distant primate species, whose lineages diverged ~29 million years ago (Kumar et al. 2017). Treatments that alter the epigenetic age of vervets according to our human-vervet clocks are likely to exert similar effects in humans.

Beyond the laboratory, vervet monkeys from wild populations are increasingly used in biomedical and anthropological research (Turner et al. 2019). Tooth eruption patterns are typically used as a practical predictor of developmental stage in wild vervets and other NHPs (Ockerse 1959; Turner et al. 1997). Although these patterns approximate the developmental stage of an individual, their utility is limited in terms of accurate prediction of chronological age, particularly in adults with a fully developed dental pattern, in which distinguishing among various stages of adulthood and senescence is difficult. In animals of unknown chronological age, the epigenetic clock can enable more accurate estimates of chronological age and thus decrease the confounding effects of age, increase the statistical power of analysis, and decrease the number of animals needed for studies, according to the ‘Three Rs’: replacement, reduction and refinement. In addition, it can improve monitoring of health status in natural populations, provide insight into life history, and enable identification of lifespan modulating factors in the context of a natural habitat.

## Materials and Methods

### Study subjects

All animals used in this study were Caribbean-origin vervet monkeys (*Chlorocebus sabaeus*) from the VRC at Wake Forest School of Medicine. The VRC colony is an extended multigenerational pedigree established from 57 founders imported from the islands of St. Kitts and Nevis in the West Indies. The introduction of new animals to the pedigree ended in the mid-1980s (Jasinska et al. 2013). The colony members are socially reared in extended family groups mimicking the natural social composition of vervet monkey troops in the wild. Group sizes range from 11 to 23 animals, with one or two intact adult males included in each group. Unfamiliar males are rotated into each group every 3–5 years. The pedigree structure is genetically confirmed (Huang et al. 2015). All colony-born vervets have known chronological age accurate to 1 day.

Beyond applications in aging studies, animals from VRC are used in a wide range of research in areas such as the efficacy and enhancement of vaccines for infectious diseases, e.g., influenza and dengue (Kim et al. 2015; Holbrook et al. 2016; Briggs et al. 2014), investigations of diabetes, metabolic disease and obesity (Kavanagh et al. 2017; Kavanagh et al. 2016; Kavanagh et al. 2013); and the development of novel non-invasive biomedical imaging methodologies (Prabhakaran et al. 2017; Maldjian et al. 2014).

### Ethics statement

The Wake Forest School of Medicine facilities are certified by the Association for Assessment and Accreditation of Laboratory Animal Care. The animal handling and sample collection procedures in this study were performed by a veterinarian after review and approval by the UCLA and VA Institutional Animal Care and Use Committees. Both housing and sample collection were in compliance with the US National Research Council Committee’s Guidelines for Care and Use of Laboratory Animals (National Research Council et al. 2011) and met or exceeded all standards of the Public Health Service’s “Policy on the Humane Care and Use of Laboratory Animals” (Office of Laboratory Animal Welfare n.d.).

### Vervet tissue samples

For this study, we selected a total of 240 samples representing the entire vervet lifespan, from neonatal to senile stages: 144 samples from the peripheral blood, 48 samples from the liver, and 48 samples from the cortical brain area BA10. The brains were perfused to remove blood prior to dissection (Jasinska et al. 2017). The targeted brain area BA10 was very small, and brain samples were dissected as bulk tissues, collecting, to the extent feasible without the benefit of microscopy, the full thickness of the cortex while avoiding the underlying white matter (Jasinska et al. 2017). One outlier blood sample (202943350003_R03C01 from animal 1992020) was excluded from analysis on the basis of the DNAm profile. The remaining 143 blood samples included 14 pairs of biological replicates collected from 14 individuals at two different time points 3.9–10.93 years apart. Peripheral blood was collected through venipuncture with standard procedures. Liver and brain cortical tissues were collected during necropsies (Jasinska et al. 2017).

Genomic DNA was isolated from blood and liver samples primarily through Puregene chemistry (Qiagen). DNA from the liver was extracted manually and that from the blood was extracted with an automated Autopure LS system (Qiagen). DNA was extracted from old liver tissues and clotted blood samples manually with a QIAamp DNA Blood Midi Kit and DNeasy Tissue Kit according to the manufacturer’s protocol (Qiagen, Valencia, CA). DNA from BA10 was extracted on an automated nucleic acid extraction platform AnaPrep (Biochain) with a magnetic bead based extraction method and Tissue DNA Extraction Kit (AnaPrep).

### Human tissue samples

To build the human-vervet clock, we analyzed previously generated methylation data from n=852 human tissue samples (adipose, blood, bone marrow, dermis, epidermis, heart, keratinocytes, fibroblasts, kidney, liver, lung, lymph node, muscle, pituitary, skin, spleen) from individuals whose ages ranged from 0 to 93 years. These human methylation data are described in (Horvath, Singh, et al. 2020). The tissue samples came from three sources. Tissue and organ samples were from the National NeuroAIDS Tissue Consortium (Morgello et al. 2001). Blood samples were from the Cape Town Adolescent Antiretroviral Cohort study (Horvath, Stein, et al. 2018). Skin and other primary cells were provided by Kenneth Raj (Kabacik et al. 2018). Ethics approval (IRB#15-001454, IRB#16-000471, IRB#18-000315, IRB#16-002028).

### Rhesus tissue samples

To validate the vervet clock cross species, we utilized the CpG methylation data described in a companion paper (Horvath, Zoller, et al. 2020).

### DNA methylation data

All DNAm data used was generated using the custom Illumina chip “HorvathMammalMethylChip40”, so called the mammalian methylation array. The mammalian methylation array is an attractive tool for DNAm assessment in primates, because it comprises 38K probes, including nearly ~36 K probes targeting CpG sites in highly conserved regions in mammals. Briefly, two thousands out of 38 K probes were selected based on their utility for human biomarker studies: these CpGs, which were previously implemented in human Illumina Infinium arrays (EPIC, 450K) were selected due to their relevance for estimating age, blood cell counts, or the proportion of neurons in brain tissue. The remaining 35,988 probes were chosen to assess cytosine DNA methylation levels in mammalian species. Toward this end, highly conserved CpGs across 50 mammalian species were selected: 33,493 Infinium II probes and 2,496 Infinium I probes. Not all probes on the array are expected to work for all species, but rather each probe is designed to cover a certain subset of species, such that overall all species have a high number of probes. The particular subset of species for each probe is provided in the chip manifest file can be found at Gene Expression Omnibus (GEO) at NCBI as platform GPL28271. The SeSaMe normalization method was used to define beta values for each probe (Zhou et al. 2018).

### Penalized Regression models

Details on the clocks (CpGs, genome coordinates) and R software code are provided in the Supplement. Penalized regression models were created with glmnet (Friedman et al. 2010). We investigated models produced by both “elastic net” regression (alpha=0.5). The optimal penalty parameters in all cases were determined automatically by using a 10 fold internal cross-validation (cv.glmnet) on the training set. By definition, the alpha value for the elastic net regression was set to 0.5 (midpoint between Ridge and Lasso type regression) and was not optimized for model performance.

We performed a cross-validation scheme for arriving at unbiased (or at least less biased) estimates of the accuracy of the different DNAm based age estimators. For validation of the clocks, we used leave-one-out LOO cross-validation (LOOCV) in which one sample was left out of the regression, then predicted the age for the remaining samples and iterated this process over all samples.

A critical step is the transformation of chronological age (the dependent variable). While no transformation was used for the multi-tissue clock for vervets, we did use a log linear transformation for the dual species clock of chronological age (Supplement).

### Relative age estimation

To introduce biological meaning into age estimates of vervets and humans that have very different lifespan; as well as to overcome the inevitable skewing due to unequal distribution of data points from vervets and humans across age range, relative age estimation was made using the formula: Relative age= Age/maxLifespan where the maximum lifespan for the two species was chosen from the *anAge* database (Magalhães et al. 2007). Maximum age of vervets and humans was 30.8 and 122.5 years, respectively.

### Epigenome wide association studies of age

EWAS was performed in each tissue separately with the R function “standardScreeningNumericTrait” in the “WGCNA” R package (Langfelder & Horvath 2008). Next the results were combined across tissues with Stouffer’s meta analysis method.

### CpG set Enrichment analysis

The significant CpGs for each tissue were selected for enrichment analysis. The first enrichment analysis was done for transcriptional factor motifs. Using the MEME motif discovery algorithm (Bailey et al. 2009), we predicted the probes that are located on TF motifs from five databases: Jasper, Taipale, Taipaledimer, Uniprob, and TRANSFAC. The overlap of selected CpGs based on the EWAS was tested with the predicted background using a hypergeometric test.

### Genome annotation

The gene-level enrichment was done using GREAT analysis (McLean et al. 2010) and human Hg19 background. The background probes were limited to 24,799 probes that were mapped to the same gene in the Vervet Monkey genome. Gene set enrichment was done for gene ontology, molecular pathways, diseases, upstream regulators, human and mouse phenotypes.

We aligned microarray probes to the vervet reference genome Chlorocebus_sabeus 1.1 GCF_000409795.2 (Warren et al. 2015). CpG sites were annotated in relation to the nearest genes based on the vervet gene annotations: Ensembl *Chlorocebus sabaeus* Annotation Release 100 (Pruitt et al. 2012). In total, 35,898 probes from the mammalian BeadChip array could be aligned to ChlSab1.1.100 genome (Ensembl). The alignment was done using the QUASR package (Gaidatzis et al. 2015), with the assumption for bisulfite conversion treatment of the genomic DNA. Following the alignment, the CpGs were annotated based on the distance to the closest transcriptional start site using the Chipseeker package (Yu et al. 2015).

## URLs

Vervet reference genome, ftp://ftp.ensembl.org/pub/release-100/fasta/chlorocebus_sabaeus/ NCBI’s vervet gene annotations, ftp://ftp.ensembl.org/pub/release-100/gtf/chlorocebus_sabaeus/

## Acknowledgements

This work was supported by the Paul G. Allen Frontiers Group (PI Steve Horvath) and the following grants from the US National Institutes of Health: P40RR019963/OD010965 (to M.J.J.); R01RR016300/OD010980 (to N.B.F.); R37MH060233 (to D. Geschwind). The rhesus macaque data were funded in part by the Intramural Research Program, National Institute on Aging, NIH

## Author contributions

KK, MJJ, RW, and NBF contributed the vervet tissue samples.

JAM contributed the rhesus data.

SH, CZL, AA, and JE developed the custom methylation array HorvathMammalMethylChip40.

SH, JDY, OW, XL, AWR, KW, and GC prepared the samples and generated the data.

AH, JZ, AJJ, and SH carried out the statistical analysis. AJJ, AH and SH designed the experiments and wrote the paper. SH conceived the study.

## SUPPLEMENTARY MATERIAL

**Supplementary Figure 1.**
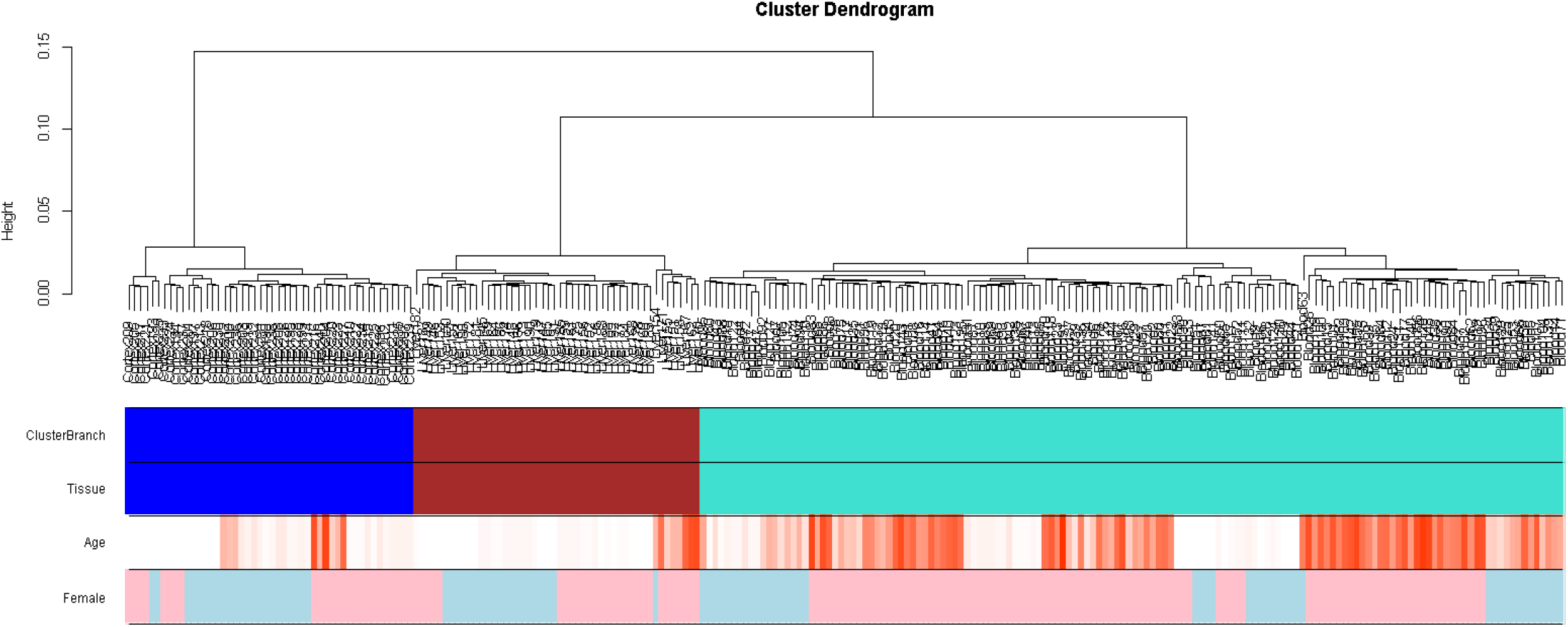
Unsupervised hierarchical clustering of tissue samples. Average linkage hierarchical clustering based on the interarray correlation coefficient (Pearson correlation). A height cut-off of 0.05 led to branch colors that correspond to Tissue type (second color band: blue - brain cortex from day 0 to 22 years of age, maroon – liver from day 0 to 21 years of age, turquoise – blood from day 1 to 25 years of age), sex (females – pink, males – blue). Age is shown in the scale from youngest (white) to oldest (red).

**Supplementary Figure 2.**
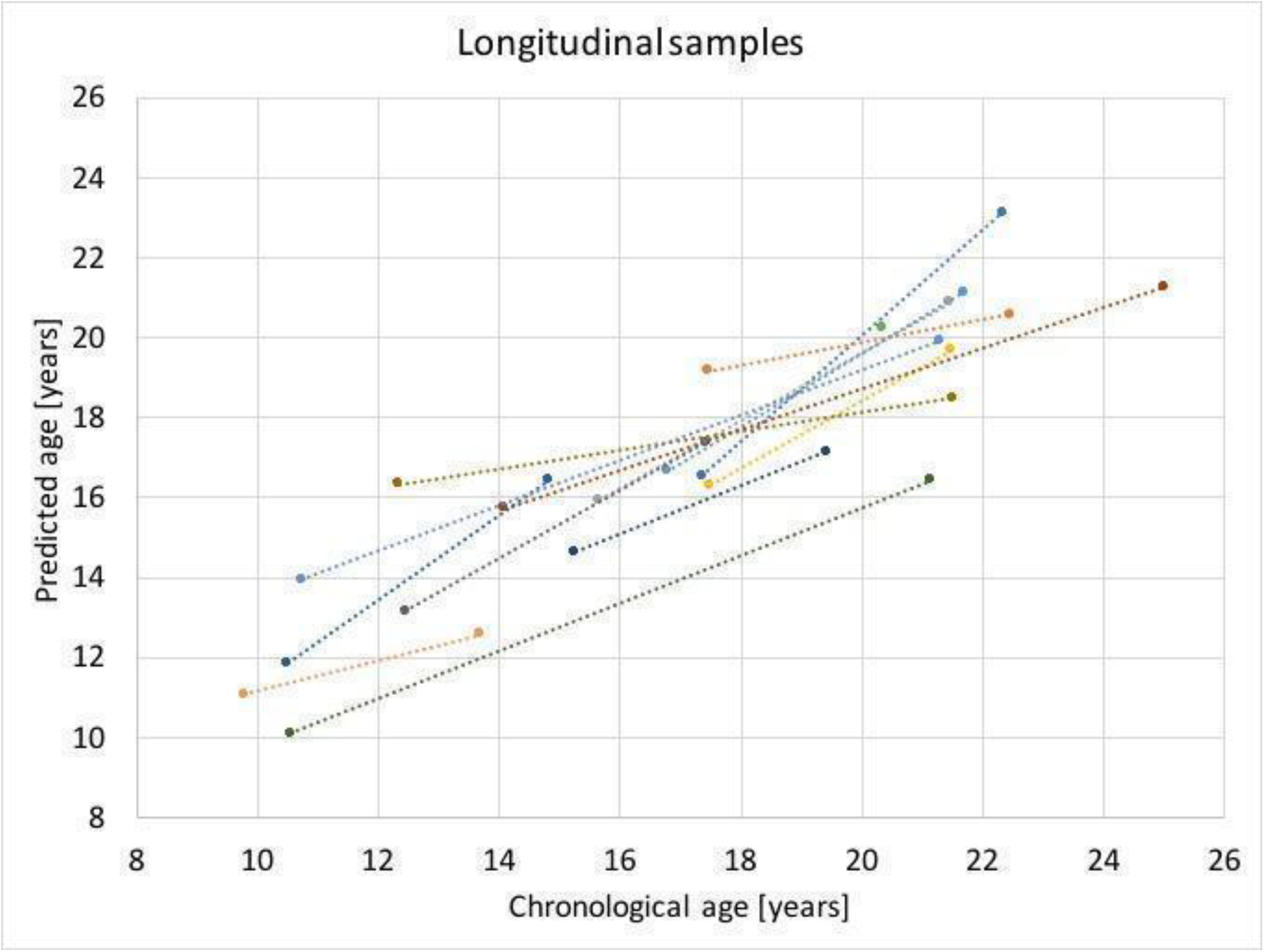
The DNAm age predictions by using the blood-specific clock in pairs of blood samples collected from the same animals at two different time points. Predicted age (y-axis) is shown relative to chronological age (x-axis) for pairs of samples collected from 14 individuals.

**Supplementary Figure 3.**
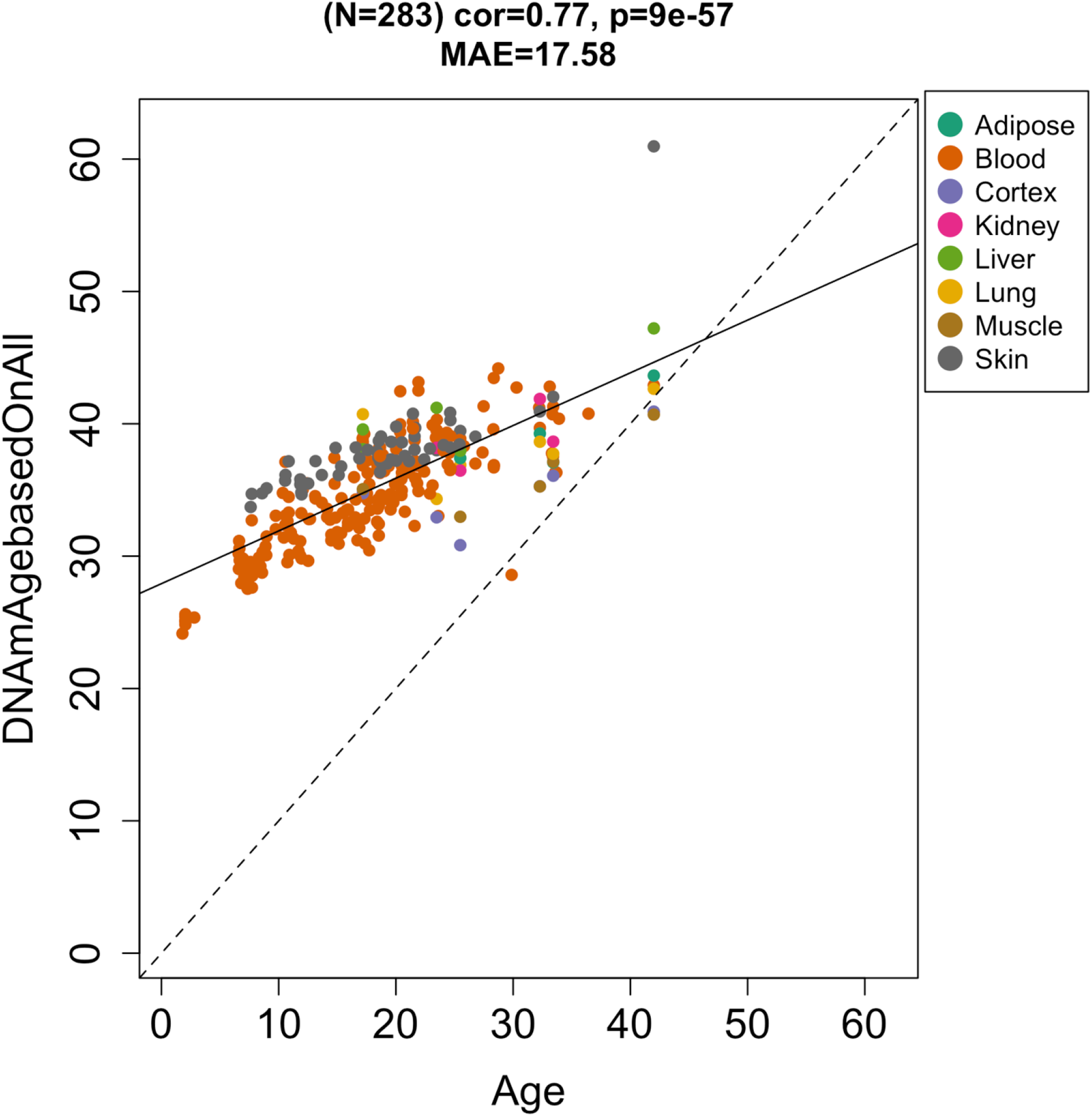
Vervet pan-clock applied to macaque tissues. The predicted DNAm age in human tissues according to the vervet pan-clock (y-axis) and chronological age of the human specimens (x-axis). The number of samples is shown in parentheses; cor – Pearson’s correlation, MAE – median absolute error.

**Supplementary Figure 4.**
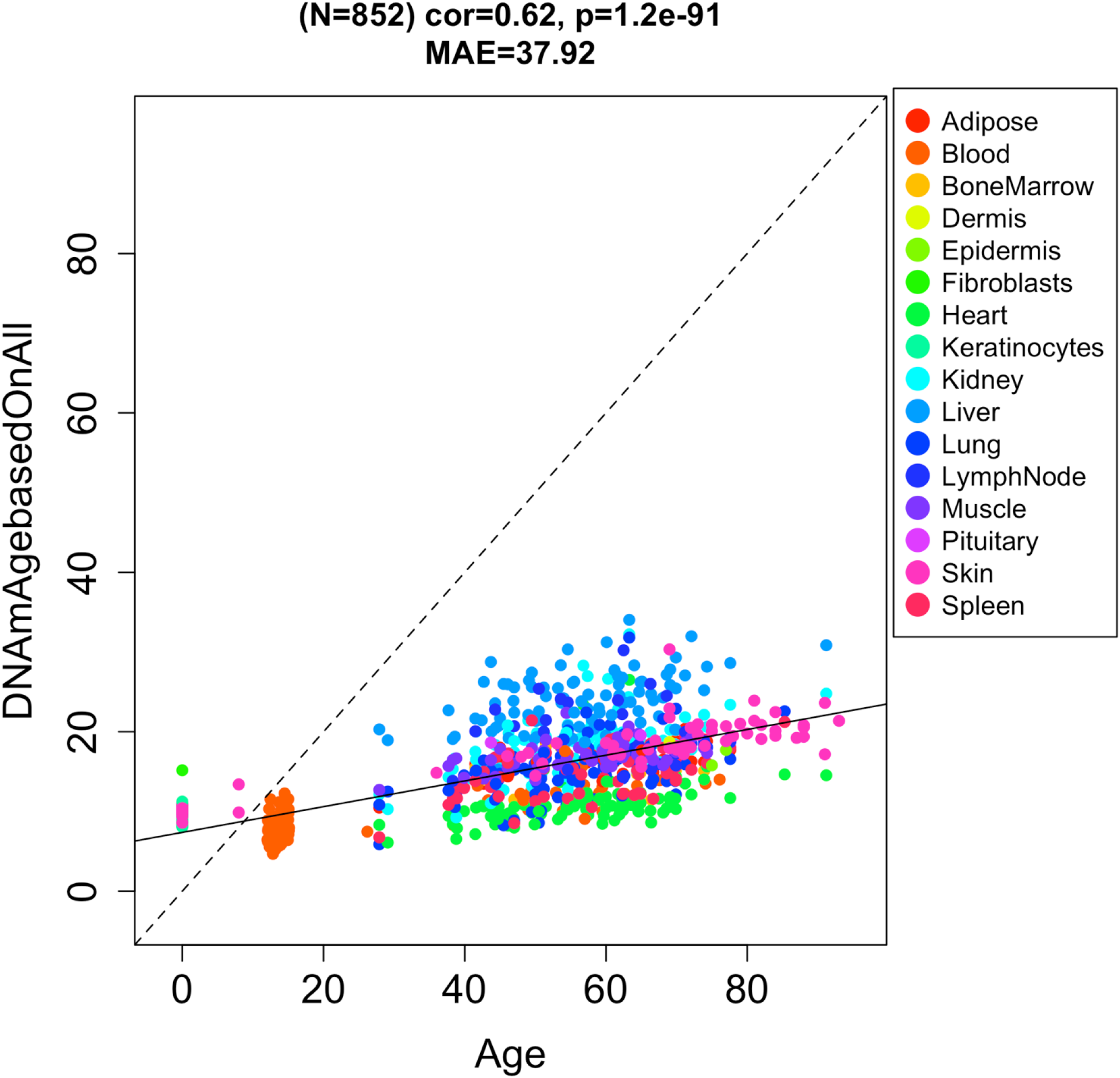
The conservation DNAm age estimated with the vervet pan-clock in human tissues. The predicted DNAm age in human tissues according to the vervet pan-clock (y-axis) and chronological age of the human specimens (x-axis). The number of samples is shown in parentheses; cor – Pearson’s correlation, MAE – median absolute error.

**Supplementary Figure 5.**
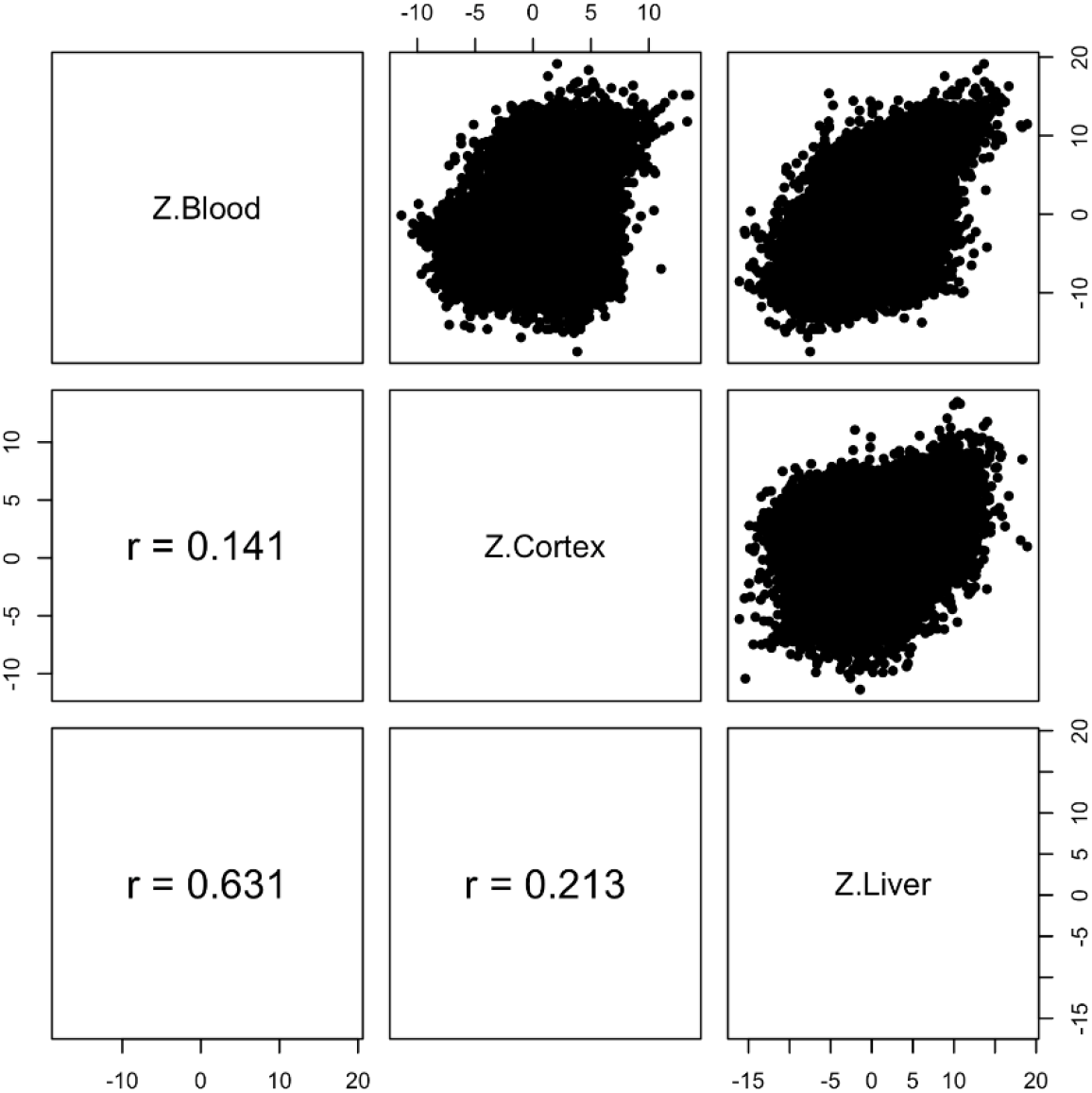
Epigenome wide association study of correlation in three different tissues. Each dot corresponds to a CpG. Z statistics for a correlation test of age in the blood, cerebral cortex, and liver. Pairwise scatter plots reveal a strong positive correlation (r=0.63) between EWAS results in the blood and brain tissue.

**Supplementary Figure 6.**
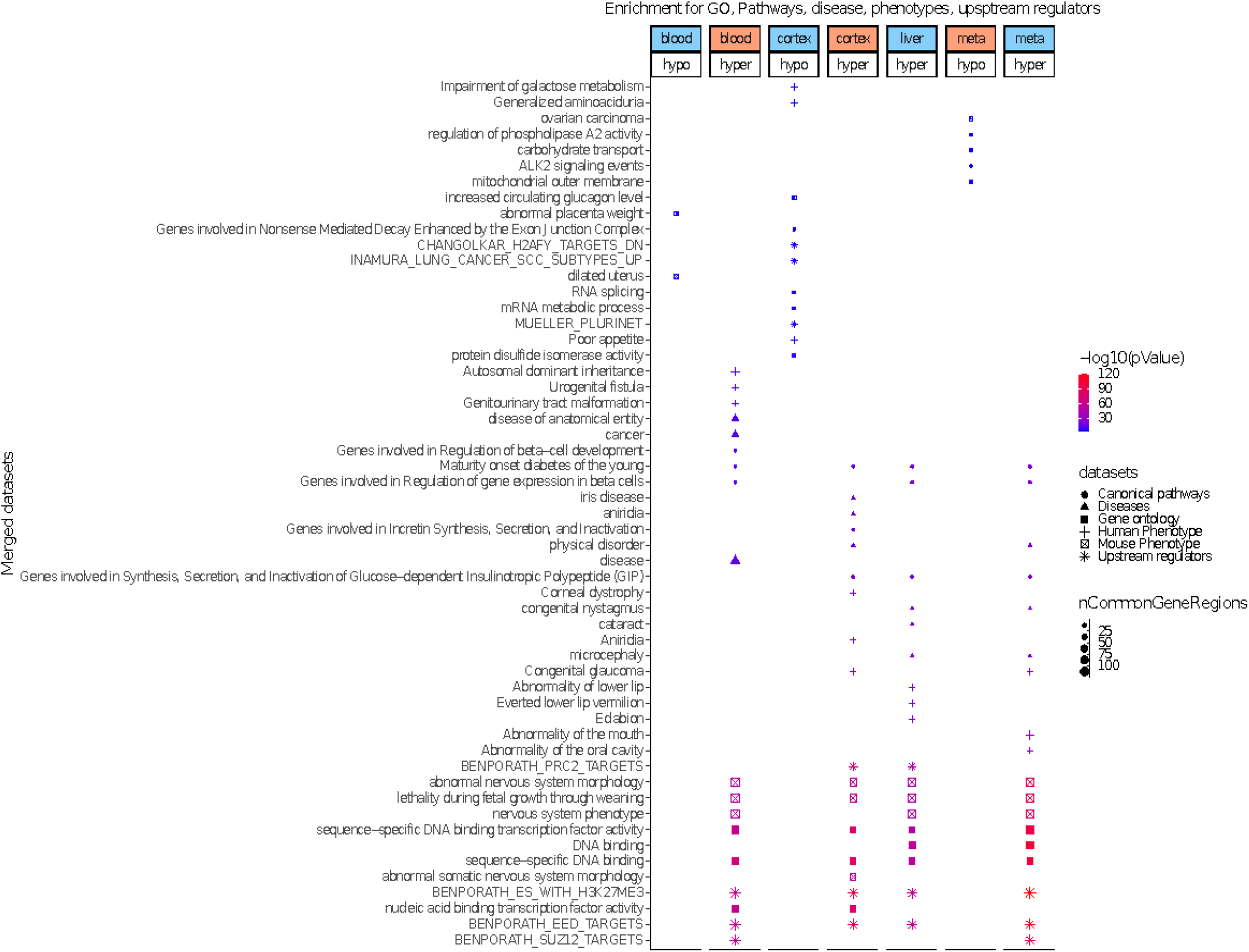
Enrichment analysis of the top CpGs associated with DNAm age in each tissue and in meta analysis. The analysis was done using the genomic region of enrichment annotation tool (McLean et al. 2010). The gene level enrichment was done using GREAT analysis (McLean et al. 2010) and human Hg19 background. The background probes were limited to 24,799 probes that were mapped to the same gene in the vervet monkey genome. The top three enriched datasets from each category (Canonical pathways, diseases, gene ontology, human and mouse phenotypes, and upstream regulators) were selected and further filtered for significance at p < 10^−4^.

## References

Ariani F, Hayek G, Rondinella D, Artuso R, Mencarelli MA, Spanhol-Rosseto A, Pollazzon M, Buoni S, Spiga O, Ricciardi S, Meloni I, Longo I, Mari F, Broccoli V, Zappella M & Renieri A (2008) FOXG1 is responsible for the congenital variant of Rett syndrome. Am. J. Hum. Genet. 83, 89–93.

Astrinidis A, Kim J, Kelly CM, Olofsson BA, Torabi B, Sorokina EM & Azizkhan-Clifford J (2010) The transcription factor SP1 regulates centriole function and chromosomal stability through a functional interaction with the mammalian target of rapamycin/raptor complex. Genes Chromosomes Cancer 49, 282–297.

Atkins HM, Willson CJ, Silverstein M, Jorgensen M, Floyd E, Kaplan JR & Appt SE (2014) Characterization of ovarian aging and reproductive senescence in vervet monkeys (Chlorocebus aethiops sabaeus). Comp. Med. 64, 55–62.

Bacalini MG, Franceschi C, Gentilini D, Ravaioli F, Zhou X, Remondini D, Pirazzini C, Giuliani C, Marasco E, Gensous N, Di Blasio AM, Ellis E, Gramignoli R, Castellani G, Capri M, Strom S, Nardini C, Cescon M, Grazi GL & Garagnani P (2019) Molecular Aging of Human Liver: An Epigenetic/Transcriptomic Signature. J. Gerontol. A Biol. Sci. Med. Sci. 74, 1–8.

Bailey TL, Boden M, Buske FA, Frith M, Grant CE, Clementi L, Ren J, Li WW & Noble WS (2009) MEME SUITE: tools for motif discovery and searching. Nucleic Acids Res. 37, W202–8.

Bartke A (2019) Growth Hormone and Aging: Updated Review. World J. Mens Health 37, 19–30.

Bell CG, Lowe R, Adams PD, Baccarelli AA, Beck S, Bell JT, Christensen BC, Gladyshev VN, Heijmans BT, Horvath S, Ideker T, Issa J-PJ, Kelsey KT, Marioni RE, Reik W, Relton CL, Schalkwyk LC, Teschendorff AE, Wagner W, Zhang K & Rakyan VK (2019) DNA methylation aging clocks: challenges and recommendations. Genome Biol. 20, 249.

Bhatlekar S, Fields JZ & Boman BM (2018) Role of HOX Genes in Stem Cell Differentiation and Cancer. Stem Cells Int. 2018, 3569493.

Bjornson-Hooper ZB, Fragiadakis GK, Spitzer MH, Madhireddy D, McIlwain D & Nolan GP (2019) A comprehensive atlas of immunological differences between humans, mice and non-human primates. bioRxiv, 574160. Available at: https://www.biorxiv.org/content/10.1101/574160v1.abstract [Accessed April 8, 2019].

Briggs CM, Smith KM, Piper A, Huitt E, Spears CJ, Quiles M, Ribeiro M, Thomas ME, Brown DT & Hernandez R (2014) Live attenuated tetravalent dengue virus host range vaccine is immunogenic in African green monkeys following a single vaccination. J. Virol. 88, 6729–6742.

Bryant A, Palma CA, Jayaswal V, Yang YW, Lutherborrow M & Ma DD (2012) miR-10a is aberrantly overexpressed in Nucleophosmin1 mutated acute myeloid leukaemia and its suppression induces cell death. Mol. Cancer 11, 8.

Chahroudi A, Bosinger SE, Vanderford TH, Paiardini M & Silvestri G (2012) Natural SIV hosts: showing AIDS the door. Science 335, 1188–1193.

Chatterjee HJ, Ho SYW, Barnes I & Groves C (2009) Estimating the phylogeny and divergence times of primates using a supermatrix approach. BMC Evol. Biol. 9, 259.

Chen BH, Marioni RE, Colicino E, Peters MJ, Ward-Caviness CK, Tsai P-C, Roetker NS, Just AC, Demerath EW, Guan W, Bressler J, Fornage M, Studenski S, Vandiver AR, Moore AZ, Tanaka T, Kiel DP, Liang L, Vokonas P, Schwartz J, Lunetta KL, Murabito JM, Bandinelli S, Hernandez DG, Melzer D, Nalls M, Pilling LC, Price TR, Singleton AB, Gieger C, Holle R, Kretschmer A, Kronenberg F, Kunze S, Linseisen J, Meisinger C, Rathmann W, Waldenberger M, Visscher PM, Shah S, Wray NR, McRae AF, Franco OH, Hofman A, Uitterlinden AG, Absher D, Assimes T, Levine ME, Lu AT, Tsao PS, Hou L, Manson JE, Carty CL, LaCroix AZ, Reiner AP, Spector TD, Feinberg AP, Levy D, Baccarelli A, van Meurs J, Bell JT, Peters A, Deary IJ, Pankow JS, Ferrucci L & Horvath S (2016) DNA methylation-based measures of biological age: meta-analysis predicting time to death. Aging 8, 1844–1865.

Chen JA, Fears SC, Jasinska AJ, Huang A, Al-Sharif NB, Scheibel KE, Dyer TD, Fagan AM, Blangero J, Woods R, Jorgensen MJ, Kaplan JR, Freimer NB & Coppola G (2018) Neurodegenerative disease biomarkers Aβ 1-40, Aβ 1-42, tau, and p-tau 181 in the vervet monkey cerebrospinal fluid: Relation to normal aging, genetic influences, and cerebral amyloid angiopathy. Brain Behav. 8, e00903.

Chen-Plotkin AS, Sadri-Vakili G, Yohrling GJ, Braveman MW, Benn CL, Glajch KE, DiRocco DP, Farrell LA, Krainc D, Gines S, MacDonald ME & Cha J-HJ (2006) Decreased association of the transcription factor Sp1 with genes downregulated in Huntington’s disease. Neurobiol. Dis. 22, 233–241.

Cole JJ, Robertson NA, Rather MI, Thomson JP, McBryan T, Sproul D, Wang T, Brock C, Clark W, Ideker T, Meehan RR, Miller RA, Brown-Borg HM & Adams PD (2017) Diverse interventions that extend mouse lifespan suppress shared age-associated epigenetic changes at critical gene regulatory regions. Genome Biol. 18, 58.

Colman RJ (2018) Non-human primates as a model for aging. Biochim. Biophys. Acta Mol. Basis Dis. 1864, 2733–2741.

Dunzendorfer S, Lee H-K & Tobias PS (2004) Flow-dependent regulation of endothelial Toll-like receptor 2 expression through inhibition of SP1 activity. Circ. Res. 95, 684–691.

Estes JD, Wong SW & Brenchley JM (2018) Nonhuman primate models of human viral infections. Nat. Rev. Immunol. 18, 390–404.

Fahy GM, Brooke RT, Watson JP, Good Z, Vasanawala SS, Maecker H, Leipold MD, Lin DTS, Kobor MS & Horvath S (2019) Reversal of epigenetic aging and immunosenescent trends in humans. Aging Cell 18, e13028.

Fairbanks LA, Bailey JN, Breidenthal SE, Laudenslager ML, Kaplan JR & Jorgensen MJ (2011) Environmental stress alters genetic regulation of novelty seeking in vervet monkeys. Genes Brain Behav. 10, 683–688.

Fairbanks LA, Jorgensen MJ, Bailey JN, Breidenthal SE, Grzywa R & Laudenslager ML (2011) Heritability and genetic correlation of hair cortisol in vervet monkeys in low and higher stress environments. Psychoneuroendocrinology 36, 1201–1208.

Finch CE & Austad SN (2012) Primate aging in the mammalian scheme: the puzzle of extreme variation in brain aging. Age 34, 1075–1091.

Finotti A, Bianchi N, Fabbri E, Borgatti M, Breveglieri G, Gasparello J & Gambari R (2015) Erythroid induction of K562 cells treated with mithramycin is associated with inhibition of raptor gene transcription and mammalian target of rapamycin complex 1 (mTORC1) functions. Pharmacological Research 91, 57–68. Available at: http://dx.doi.org/10.1016/j.phrs.2014.11.005.

Friedman J, Hastie T & Tibshirani R (2010) Regularization Paths for Generalized Linear Models via Coordinate Descent. J. Stat. Softw. 33, 1–22.

Gaidatzis D, Lerch A, Hahne F & Stadler MB (2015) QuasR: quantification and annotation of short reads in R. Bioinformatics 31, 1130–1132.

Gibson J, Russ TC, Clarke T-K, Howard DM, Hillary RF, Evans KL, Walker RM, Bermingham ML, Morris SW, Campbell A, Hayward C, Murray AD, Porteous DJ, Horvath S, Lu AT, McIntosh AM, Whalley HC & Marioni RE (2019) A meta-analysis of genome-wide association studies of epigenetic age acceleration. PLoS Genet. 15, e1008104.

Hatzipetros T, Bogdanik LP, Tassinari VR, Kidd JD, Moreno AJ, Davis C, Osborne M, Austin A, Vieira FG, Lutz C & Perrin S (2014) C57BL/6J congenic Prp-TDP43A315T mice develop progressive neurodegeneration in the myenteric plexus of the colon without exhibiting key features of ALS. Brain Res. 1584, 59–72.

Holbrook BC, Kim JR, Blevins LK, Jorgensen MJ, Kock ND, D’Agostino RBJr, Aycock ST, Hadimani MB, King SB, Parks GD & Alexander-Miller MA (2016) A Novel R848-Conjugated Inactivated Influenza Virus Vaccine Is Efficacious and Safe in a Neonate Nonhuman Primate Model. J. Immunol. 197, 555–564.

Horvath S (2013) DNA methylation age of human tissues and cell types. Genome Biol. 14, R115.

Horvath S & Levine AJ (2015) HIV-1 Infection Accelerates Age According to the Epigenetic Clock. J. Infect. Dis. 212, 1563–1573.

Horvath S, Oshima J, Martin GM, Lu AT, Quach A, Cohen H, Felton S, Matsuyama M, Lowe D, Kabacik S, Wilson JG, Reiner AP, Maierhofer A, Flunkert J, Aviv A, Hou L, Baccarelli AA, Li Y, Stewart JD, Whitsel EA, Ferrucci L, Matsuyama S & Raj K (2018) Epigenetic clock for skin and blood cells applied to Hutchinson Gilford Progeria Syndrome and ex vivo studies. Aging 10, 1758–1775.

Horvath S & Raj K (2018) DNA methylation-based biomarkers and the epigenetic clock theory of ageing. Nat. Rev. Genet. 19, 371–384.

Horvath S, Singh K, Raj K, Khairnar S & Sanghavi A (2020) Reversing age: dual species measurement of epigenetic age with a single clock. bioRxiv. Available at: https://www.biorxiv.org/content/10.1101/2020.05.07.082917v1.

Horvath S, Stein DJ, Phillips N, Heany SJ, Kobor MS, Lin DTS, Myer L, Zar HJ, Levine AJ & Hoare J (2018) Perinatally acquired HIV infection accelerates epigenetic aging in South African adolescents. AIDS 32, 1465–1474.

Horvath S, Zoller JA, Haghani A, Lu AT, Li CZ, Raj K, Jasinska AJ & Mattison JA (2020) Epigenetic clock and methylation studies in rhesus macaque.

Huang YS, Ramensky V, Service SK, Jasinska AJ, Jung Y, Choi O-W, Cantor RM, Juretic N, Wasserscheid J, Kaplan JR, Jorgensen MJ, Dyer TD, Dewar K, Blangero J, Wilson RK, Warren W, Weinstock GM & Freimer NB (2015) Sequencing strategies and characterization of 721 vervet monkey genomes for future genetic analyses of medically relevant traits. BMC Biol. 13, 41.

Jasinska AJ (2019) Biological Resources for Genomic Investigation in the Vervet Monkey (Chlorocebus). Savanna Monkeys, 16–28. Available at: http://dx.doi.org/10.1017/9781139019941.002.

Jasinska AJ, Lin MK, Service S, Choi O-W, DeYoung J, Grujic O, Kong S-Y, Jung Y, Jorgensen MJ, Fairbanks LA, Turner T, Cantor RM, Wasserscheid J, Dewar K, Warren W, Wilson RK, Weinstock G, Jentsch JD & Freimer NB (2012) A non-human primate system for large-scale genetic studies of complex traits. Hum. Mol. Genet. 21, 3307–3316.

Jasinska AJ, Pandrea I, He T, Benjamin C, Newton M, Lee JC, Freimer NB, Coppola G & Jentsch JD (2020) Immunosuppressive effect and global dysregulation of blood transcriptome in response to psychosocial stress in vervet monkeys (Chlorocebus sabaeus). Scientific Reports 10. Available at: http://dx.doi.org/10.1038/s41598-020-59934-z.

Jasinska AJ, Rostamian D, Davis AT & Kavanagh K (2020) Transcriptomic Analysis of Cell-free Fetal RNA in the Amniotic Fluid of Vervet Monkeys (Chlorocebus sabaeus). Comp. Med. 70, 67–74.

Jasinska AJ, Schmitt CA, Service SK, Cantor RM, Dewar K, Jentsch JD, Kaplan JR, Turner TR, Warren WC, Weinstock GM, Woods RP & Freimer NB (2013) Systems biology of the vervet monkey. ILAR J. 54, 122–143.

Jasinska AJ, Service S, Choi O-W, DeYoung J, Grujic O, Kong S-Y, Jorgensen MJ, Bailey J, Breidenthal S, Fairbanks LA, Woods RP, Jentsch JD & Freimer NB (2009) Identification of brain transcriptional variation reproduced in peripheral blood: an approach for mapping brain expression traits. Hum. Mol. Genet. 18, 4415–4427.

Jasinska AJ, Zelaya I, Service SK, Peterson CB, Cantor RM, Choi O-W, DeYoung J, Eskin E, Fairbanks LA, Fears S, Furterer AE, Huang YS, Ramensky V, Schmitt CA, Svardal H, Jorgensen MJ, Kaplan JR, Villar D, Aken BL, Flicek P, Nag R, Wong ES, Blangero J, Dyer TD, Bogomolov M, Benjamini Y, Weinstock GM, Dewar K, Sabatti C, Wilson RK, Jentsch JD, Warren W, Coppola G, Woods RP & Freimer NB (2017) Genetic variation and gene expression across multiple tissues and developmental stages in a nonhuman primate. Nat. Genet. 49, 1714–1721.

Johnson AA, Akman K, Calimport SRG, Wuttke D, Stolzing A & de Magalhães JP (2012) The role of DNA methylation in aging, rejuvenation, and age-related disease. Rejuvenation Res. 15, 483–494.

Kabacik S, Horvath S, Cohen H & Raj K (2018) Epigenetic ageing is distinct from senescence-mediated ageing and is not prevented by telomerase expression. Aging 10, 2800–2815.

Kalinin S, Willard SL, Shively CA, Kaplan JR, Register TC, Jorgensen MJ, Polak PE, Rubinstein I & Feinstein DL (2013) Development of amyloid burden in African Green monkeys. Neurobiol. Aging 34, 2361–2369.

Kavanagh K, Davis AT, Jenkins KA & Flynn DM (2016) Effects of heated hydrotherapy on muscle HSP70 and glucose metabolism in old and young vervet monkeys. Cell Stress Chaperones 21, 717–725.

Kavanagh K, Davis AT, Peters DE, LeGrand AC, Bharadwaj MS & Molina AJA (2017) Regulators of mitochondrial quality control differ in subcutaneous fat of metabolically healthy and unhealthy obese monkeys. Obesity 25, 689–696.

Kavanagh K, Wylie AT, Tucker KL, Hamp TJ, Gharaibeh RZ, Fodor AA & Cullen JMC (2013) Dietary fructose induces endotoxemia and hepatic injury in calorically controlled primates. Am. J. Clin. Nutr. 98, 349–357.

Kennerdell JR, Liu N & Bonini NM (2018) MiR-34 inhibits polycomb repressive complex 2 to modulate chaperone expression and promote healthy brain aging. Nature Communications 9. Available at: http://dx.doi.org/10.1038/s41467-018-06592-5.

Kim JR, Holbrook BC, Hayward SL, Blevins LK, Jorgensen MJ, Kock ND, De Paris K, D’Agostino RBJr, Aycock ST, Mizel SB, Parks GD & Alexander-Miller MA (2015) Inclusion of Flagellin during Vaccination against Influenza Enhances Recall Responses in Nonhuman Primate Neonates. J. Virol. 89, 7291–7303.

Kondo T, Sheets PL, Zopf DA, Aloor HL, Cummins TR, Chan RJ & Hashino E (2008) Tlx3 exerts context-dependent transcriptional regulation and promotes neuronal differentiation from embryonic stem cells. Proc. Natl. Acad. Sci. U. S. A. 105, 5780–5785.

Kresovich JK, Xu Z, O’Brien KM, Weinberg CR, Sandler DP & Taylor JA (2019) Methylation-Based Biological Age and Breast Cancer Risk. J. Natl. Cancer Inst. 111, 1051–1058.

Kumar S, Stecher G, Suleski M & Hedges SB (2017) TimeTree: A Resource for Timelines, Timetrees, and Divergence Times. Mol. Biol. Evol. 34, 1812–1819.

Kuokkanen S, Polotsky AJ, Chosich J, Bradford AP, Jasinska A, Phang T, Santoro N & Appt SE (2016) Corpus luteum as a novel target of weight changes that contribute to impaired female reproductive physiology and function. Syst. Biol. Reprod. Med. 62, 227–242.

Langfelder P & Horvath S (2008) WGCNA: an R package for weighted correlation network analysis. BMC Bioinformatics9, 559.

Lankau EW, Turner PV, Mullan RJ & Galland GG (2014) Use of nonhuman primates in research in North America. J. Am. Assoc. Lab. Anim. Sci. 53, 278–282.

Latimer CS, Shively CA, Keene CD, Jorgensen MJ, Andrews RN, Register TC, Montine TJ, Wilson AM, Neth BJ, Mintz A, Maldjian JA, Whitlow CT, Kaplan JR & Craft S (2019) A nonhuman primate model of early Alzheimer’s disease pathologic change: Implications for disease pathogenesis. Alzheimers. Dement. 15, 93–105.

Levine ME, Lu AT, Chen BH, Hernandez DG, Singleton AB, Ferrucci L, Bandinelli S, Salfati E, Manson JE, Quach A, Kusters CDJ, Kuh D, Wong A, Teschendorff AE, Widschwendter M, Ritz BR, Absher D, Assimes TL & Horvath S (2016) Menopause accelerates biological aging. Proc. Natl. Acad. Sci. U. S. A. 113, 9327–9332.

Levine ME, Lu AT, Quach A, Chen BH, Assimes TL, Bandinelli S, Hou L, Baccarelli AA, Stewart JD, Li Y, Whitsel EA, Wilson JG, Reiner AP, Aviv A, Lohman K, Liu Y, Ferrucci L & Horvath S (2018) An epigenetic biomarker of aging for lifespan and healthspan. Aging 10, 573–591.

Lu AT, Hannon E, Levine ME, Hao K, Crimmins EM, Lunnon K, Kozlenkov A, Mill J, Dracheva S & Horvath S (2016) Genetic variants near MLST8 and DHX57 affect the epigenetic age of the cerebellum. Nat. Commun. 7, 10561.

Lu AT, Quach A, Wilson JG, Reiner AP, Aviv A, Raj K, Hou L, Baccarelli AA, Li Y, Stewart JD & Others (2019) DNA methylation GrimAge strongly predicts lifespan and healthspan. Aging 11, 303.

Lu AT, Xue L, Salfati EL, Chen BH, Ferrucci L, Levy D, Joehanes R, Murabito JM, Kiel DP, Tsai P-C, Yet I, Bell JT, Mangino M, Tanaka T, McRae AF, Marioni RE, Visscher PM, Wray NR, Deary IJ, Levine ME, Quach A, Assimes T, Tsao PS, Absher D, Stewart JD, Li Y, Reiner AP, Hou L, Baccarelli AA, Whitsel EA, Aviv A, Cardona A, Day FR, Wareham NJ, Perry JRB, Ong KK, Raj K, Lunetta KL & Horvath S (2018) GWAS of epigenetic aging rates in blood reveals a critical role for TERT. Nature Communications 9. Available at: http://dx.doi.org/10.1038/s41467-017-02697-5.

Ma D, Jasinska AJ, Feyertag F, Wijewardana V, Kristoff J, He T, Raehtz K, Schmitt CA, Jung Y, Cramer JD, Dione M, Antonio M, Tracy R, Turner T, Robertson DL, Pandrea I, Freimer N, Apetrei C & International Vervet Research Consortium (2014) Factors associated with siman immunodeficiency virus transmission in a natural African nonhuman primate host in the wild. J. Virol. 88, 5687–5705.

Ma D, Jasinska A, Kristoff J, Grobler JP, Turner T, Jung Y, Schmitt C, Raehtz K, Feyertag F, Martinez Sosa N, Wijewardana V, Burke DS, Robertson DL, Tracy R, Pandrea I, Freimer N, Apetrei C & International Vervet Research Consortium (2013) SIVagm infection in wild African green monkeys from South Africa: epidemiology, natural history, and evolutionary considerations. PLoS Pathog. 9, e1003011.

Magalhães JP, de Costa J & Church GM (2007) An Analysis of the Relationship Between Metabolism, Developmental Schedules, and Longevity Using Phylogenetic Independent Contrasts. J. Gerontol. A Biol. Sci. Med. Sci. 62, 149–160.

Maldjian JA, Daunais JB, Friedman DP & Whitlow CT (2014) Vervet MRI atlas and label map for fully automated morphometric analyses. Neuroinformatics 12, 543–550.

Martin RD (2003) Primatology as an essential basis for biological anthropology. Evol. Anthropol. 11, 3–6.

Mattison JA & Vaughan KL (2017) An overview of nonhuman primates in aging research. Exp. Gerontol. 94, 41–45.

McLean CY, Bristor D, Hiller M, Clarke SL, Schaar BT, Lowe CB, Wenger AM & Bejerano G (2010) GREAT improves functional interpretation of cis-regulatory regions. Nat. Biotechnol. 28, 495–501.

Meer MV, Podolskiy DI, Tyshkovskiy A & Gladyshev VN (2018) A whole lifespan mouse multi-tissue DNA methylation clock. eLife 7. Available at: http://dx.doi.org/10.7554/elife.40675.

Meyer JS & Hamel AF (2014) Models of stress in nonhuman primates and their relevance for human psychopathology and endocrine dysfunction. ILAR J. 55, 347–360.

Morgello S, Gelman BB, Kozlowski PB, Vinters HV, Masliah E, Cornford M, Cavert W, Marra C, Grant I & Singer EJ (2001) The National NeuroAIDS Tissue Consortium: a new paradigm in brain banking with an emphasis on infectious disease. Neuropathol. Appl. Neurobiol. 27, 326–335.

Mundell NA & Labosky PA (2011) Neural crest stem cell multipotency requires Foxd3 to maintain neural potential and repress mesenchymal fates. Development 138, 641–652.

National Research Council, Division on Earth and Life Studies, Institute for Laboratory AnimalResearch & Committee for the Update of the Guide for the Care and Use of Laboratory Animals (2011) Guide for the Care and Use of Laboratory Animals: Eighth Edition, National Academies Press.

Ockerse (1959) The eruption sequence and eruption times of the teeth of the vervet monkey. J. Dent. Assoc. S. Afr. 14, 422–424.

Office of Laboratory Animal Welfare PHS Policy on Humane Care and Use of Laboratory Animals | OLAW. Available at: https://olaw.nih.gov/policies-laws/phs-policy.htm [Accessed January 13, 2020].

Pandrea I, Apetrei C, Dufour J, Dillon N, Barbercheck J, Metzger M, Jacquelin B, Bohm R, Marx PA, Sinoussi F, Hirsch VM, Müller-Trutwin MC, Lackner AA & Veazey RS (2006) Simian immunodeficiency virus SIVagm.sab infection of Caribbean African green monkeys: a new model for the study of SIV pathogenesis in natural hosts. J. Virol. 80, 4858–4867.

Pasumarthy KK, Jayavelu ND, Kilpinen L, Andrus C, Battle SL, Korhonen M, Lehenkari P, Lund R, Laitinen S & David Hawkins R (2017) Methylome Analysis of Human Bone Marrow MSCs Reveals Extensive Age- and Culture-Induced Changes at Distal Regulatory Elements. Stem Cell Reports 9, 999–1015. Available at: http://dx.doi.org/10.1016/j.stemcr.2017.07.018.

Penna E, Orso F, Cimino D, Vercellino I, Grassi E, Quaglino E, Turco E & Taverna D (2013) miR-214 coordinates melanoma progression by upregulating ALCAM through TFAP2 and miR-148b downmodulation. Cancer Res. 73, 4098–4111.

Perrin S (2014) Preclinical research: Make mouse studies work. Nature 507, 423–425.

Petkovich DA, Podolskiy DI, Lobanov AV, Lee S-G, Miller RA & Gladyshev VN (2017) Using DNA Methylation Profiling to Evaluate Biological Age and Longevity Interventions. Cell Metab. 25, 954–960.e6.

Postupna N, Latimer CS, Larson EB, Sherfield E, Paladin J, Shively CA, Jorgensen MJ, Andrews RN, Kaplan JR, Crane PK, Montine KS, Craft S, Keene CD & Montine TJ (2017) Human Striatal Dopaminergic and Regional Serotonergic Synaptic Degeneration with Lewy Body Disease and Inheritance of APOE ε4. Am. J. Pathol. 187, 884–895.

Prabhakaran J, Sai KKS, Zanderigo F, Rubin-Falcone H, Jorgensen MJ, Kaplan JR, Tooke KI, Mintz A, John Mann J & Dileep Kumar JS (2017) In vivo evaluation of [18 F]FECIMBI-36, an agonist 5-HT 2A/2C receptor PET radioligand in nonhuman primate. Bioorganic & Medicinal Chemistry Letters 27, 21–23. Available at: http://dx.doi.org/10.1016/j.bmcl.2016.11.043.

Pruitt KD, Tatusova T, Brown GR & Maglott DR (2012) NCBI Reference Sequences (RefSeq): current status, new features and genome annotation policy. Nucleic Acids Res. 40, D130–5.

Quach A, Levine ME, Tanaka T, Lu AT, Chen BH, Ferrucci L, Ritz B, Bandinelli S, Neuhouser ML, Beasley JM & Others (2017) Epigenetic clock analysis of diet, exercise, education, and lifestyle factors. Aging 9, 419.

Ramensky V, Jasinska AJ, Deverasetty S, Svardal H, Zelaya I, Jorgensen MJ, Kaplan JR, Cline M, Zharikova A, Susan K. Service, Wilson RK, Coppola G, Freimer NB & Warren} WC (2019) The burden of deleterious variants in a non-human primate biomedical model. Available at: bioRxiv 784132; doi: https://doi.org/10.1101/784132.

Respuela P, Nikolić M, Tan M, Frommolt P, Zhao Y, Wysocka J & Rada-Iglesias A (2016) Foxd3 Promotes Exit from Naive Pluripotency through Enhancer Decommissioning and Inhibits Germline Specification. Cell Stem Cell 18, 118–133.

Rieger E, Bijl JJ, van Oostveen JW, Soyer HP, Oudejans CB, Jiwa NM, Walboomers JM & Meijer CJ (1994) Expression of the homeobox gene HOXC4 in keratinocytes of normal skin and epithelial skin tumors is correlated with differentiation. J. Invest. Dermatol. 103, 341–346.

Rogers J (2018) The behavioral genetics of nonhuman primates: Status and prospects. Am. J. Phys. Anthropol. 165 Suppl 65, 23–36.

Santpere G, Nieto M, Puig B & Ferrer I (2006) Abnormal Sp1 transcription factor expression in Alzheimer disease and tauopathies. Neurosci. Lett. 397, 30–34.

Schmitt CA, Bergey CM, Jasinska AJ, Ramensky V, Burt F, Svardal H, Jorgensen MJ, Freimer NB, Paul Grobler J & Turner TR (2020) ACE2 and TMPRSS2 variation in savanna monkeys (Chlorocebus spp.): Potential risk for zoonotic/anthroponotic transmission of SARS-CoV-2 and a potential model for functional studies. PLOS ONE 15, e0235106. Available at: http://dx.doi.org/10.1371/journal.pone.0235106.

Schmitt CA, Service SK, Jasinska AJ, Dyer TD, Jorgensen MJ, Cantor RM, Weinstock GM, Blangero J, Kaplan JR & Freimer NB (2018) Obesity and obesogenic growth are both highly heritable and modified by diet in a nonhuman primate model, the African green monkey (Chlorocebus aethiops sabaeus). Int. J. Obes. 42, 765–774.

Seok J, Warren HS, Cuenca AG, Mindrinos MN, Baker HV, Xu W, Richards DR, McDonald-Smith GP, Gao H, Hennessy L, Finnerty CC, López CM, Honari S, Moore EE, Minei JP, Cuschieri J, Bankey PE, Johnson JL, Sperry J, Nathens AB, Billiar TR, West MA, Jeschke MG, Klein MB, Gamelli RL, Gibran NS, Brownstein BH, Miller-Graziano C, Calvano SE, Mason PH, Cobb JP, Rahme LG, Lowry SF, Maier RV, Moldawer LL, Herndon DN, Davis RW, Xiao W, Tompkins RG & the Inflammation and Host Response to Injury LSCRP (2013) Genomic responses in mouse models poorly mimic human inflammatory diseases. Proceedings of the National Academy of Sciences 110, 3507–3512.

Stubbs TM, Bonder MJ, Stark A-K, Krueger F, BI Ageing Clock Team, von Meyenn F, Stegle O & Reik W (2017) Multi-tissue DNA methylation age predictor in mouse. Genome Biol. 18, 68.

Svardal H, Jasinska AJ, Apetrei C, Coppola G, Huang Y, Schmitt CA, Jacquelin B, Ramensky V, Müller-Trutwin M, Antonio M, Weinstock G, Grobler JP, Dewar K, Wilson RK, Turner TR, Warren WC, Freimer NB & Nordborg M (2017) Ancient hybridization and strong adaptation to viruses across African vervet monkey populations. Nat. Genet. 49, 1705–1713.

Tasic B, Menon V, Nguyen TN, Kim TK, Jarsky T, Yao Z, Levi B, Gray LT, Sorensen SA, Dolbeare T, Bertagnolli D, Goldy J, Shapovalova N, Parry S, Lee C, Smith K, Bernard A, Madisen L, Sunkin SM, Hawrylycz M, Koch C & Zeng H (2016) Adult mouse cortical cell taxonomy revealed by single cell transcriptomics. Nat. Neurosci. 19, 335–346.

Teng L, Mundell NA, Frist AY, Wang Q & Labosky PA (2008) Requirement for Foxd3 in the maintenance of neural crest progenitors. Development 135, 1615–1624.

Thompson MJ, Chwiałkowska K, Rubbi L, Lusis AJ, Davis RC, Srivastava A, Korstanje R, Churchill GA, Horvath S & Pellegrini M (2018) A multi-tissue full lifespan epigenetic clock for mice. Aging 10, 2832–2854.

Turner TR, Anapol F & Jolly CJ (1997) Growth, development, and sexual dimorphism in vervet monkeys (Cercopithecus aethiops) at four sites in Kenya. Am. J. Phys. Anthropol. 103, 19–35.

Turner TR, Schmitt CA & Cramer JD (2019) Savanna Monkeys: The Genus Chlorocebus, Cambridge University Press.

Turner TR, Schmitt CA, Cramer JD, Lorenz J, Grobler JP, Jolly CJ & Freimer NB (2018) Morphological variation in the genus Chlorocebus: Ecogeographic and anthropogenically mediated variation in body mass, postcranial morphology, and growth. Am. J. Phys. Anthropol. Available at: http://dx.doi.org/10.1002/ajpa.23459.

Vallender EJ & Miller GM (2013) Nonhuman primate models in the genomic era: a paradigm shift. ILAR J. 54, 154–165.

Voruganti VS, Jorgensen MJ, Kaplan JR, Kavanagh K, Rudel LL, Temel R, Fairbanks LA & Comuzzie AG (2013) Significant genotype by diet (G × D) interaction effects on cardiometabolic responses to a pedigree-wide, dietary challenge in vervet monkeys (Chlorocebus aethiops sabaeus). Am. J. Primatol. 75, 491–499.

Wang T, Tsui B, Kreisberg JF, Robertson NA, Gross AM, Yu MK, Carter H, Brown-Borg HM, Adams PD & Ideker T (2017) Epigenetic aging signatures in mice livers are slowed by dwarfism, calorie restriction and rapamycin treatment. Genome Biol. 18, 57.

Warren WC, Jasinska AJ, García-Pérez R, Svardal H, Tomlinson C, Rocchi M, Archidiacono N, Capozzi O, Minx P, Montague MJ, Kyung K, Hillier LW, Kremitzki M, Graves T, Chiang C, Hughes J, Tran N, Huang Y, Ramensky V, Choi O-W, Jung YJ, Schmitt CA, Juretic N, Wasserscheid J, Turner TR, Wiseman RW, Tuscher JJ, Karl JA, Schmitz JE, Zahn R, O’Connor DH, Redmond E, Nisbett A, Jacquelin B, Müller-Trutwin MC, Brenchley JM, Dione M, Antonio M, Schroth GP, Kaplan JR, Jorgensen MJ, Thomas GWC, Hahn MW, Raney BJ, Aken B, Nag R, Schmitz J, Churakov G, Noll A, Stanyon R, Webb D, Thibaud-Nissen F, Nordborg M, Marques-Bonet T, Dewar K, Weinstock GM, Wilson RK & Freimer NB (2015) The genome of the vervet (Chlorocebus aethiops sabaeus). Genome Res. 25, 1921–1933.

Xu Y, Lopes C, Qian Y, Liu Y, Cheng L, Goulding M, Turner EE, Lima D & Ma Q (2008) Tlx1 and Tlx3 coordinate specification of dorsal horn pain-modulatory peptidergic neurons. J. Neurosci. 28, 4037–4046.

Yang Z & Kaye DM (2009) Mechanistic insights into the link between a polymorphism of the 3′UTR of the SLC7A1 gene and hypertension. Hum. Mutat. 30, 328–333.

Yu G, Wang L-G & He Q-Y (2015) ChIPseeker: an R/Bioconductor package for ChIP peak annotation, comparison and visualization. Bioinformatics 31, 2382–2383.

Zeisel A, Muñoz-Manchado AB, Codeluppi S, Lönnerberg P, La Manno G, Juréus A, Marques S, Munguba H, He L, Betsholtz C, Rolny C, Castelo-Branco G, Hjerling-Leffler J & Linnarsson S (2015) Brain structure. Cell types in the mouse cortex and hippocampus revealed by single-cell RNA-seq. Science 347, 1138–1142.

Zhang J-P, Zhang H, Wang H-B, Li Y-X, Liu G-H, Xing S, Li M-Z & Zeng M-S (2014) Down-regulation of Sp1 suppresses cell proliferation, clonogenicity and the expressions of stem cell markers in nasopharyngeal carcinoma. J. Transl. Med. 12, 222.

Zheng SC, Widschwendter M & Teschendorff AE (2016) Epigenetic drift, epigenetic clocks and cancer risk. Epigenomics 8, 705–719.

Zhou W, Triche TJJr, Laird PW & Shen H (2018) SeSAMe: reducing artifactual detection of DNA methylation by Infinium BeadChips in genomic deletions. Nucleic Acids Res. 46, e123.

